# A widely-occurring family of pore-forming effectors broadens the impact of the *Serratia* Type VI secretion system

**DOI:** 10.1101/2024.11.27.625605

**Authors:** Mark Reglinski, Quenton W. Hurst, David J. Williams, Marek Gierlinski, Alp Tegin Şahin, Katharine Mathers, Adam Ostrowski, Megan Bergkessel, Ulrich Zachariae, Samantha J. Pitt, Sarah J. Coulthurst

## Abstract

The ability to compete with diverse competitors is essential for bacteria to succeed in microbial communities. A widespread strategy for inter-bacterial competition is the delivery of antibacterial toxins, or effector proteins, directly into rival cells using the Type VI secretion system (T6SS). Whilst a large number of broad-spectrum enzymatic T6SS effectors have been described, relatively few which form pores in target cell membranes have been reported. Here, we describe a widely-occurring new family of T6SS-dependent pore-forming effectors, exemplified by Ssp4 of *Serratia marcescens* Db10. We show *in vitro* that Ssp4 forms regulated pores that have higher selectivity for cations and use molecular dynamics simulations to support a high resolution structural model of a tetrameric membrane pore formed by Ssp4. Notably, Ssp4 displays a distinct ion selectivity, phylogenetic distribution and impact on intoxicated cells compared with Ssp6, the other cation-selective pore-forming toxin delivered by the same T6SS. Ssp4 is also active against a wider range of target species than Ssp6, highlighting that T6SS effectors are not always broad-spectrum. Finally, use of Tn-seq to identify Ssp4-resistant mutants reveals that a *mucA* mutant of *Pseudomonas fluorescens*, which overproduces extracellular polysaccharide, provides resistance to T6SS attacks. We conclude that possession of two distinct T6SS-dependent pore-forming toxins may be a common strategy to ensure effective de-energisation of closely- and distantly-related competitors.

## Introduction

Bacteria typically exist in mixed microbial communities, where competition for resources between and within species represents a constant challenge and a key driver for the evolution of antimicrobial toxins and the molecular machineries that deliver them to competitor cells^1, 2^. The Type VI secretion system (T6SS) occurs widely in Gram-negative bacteria and is used to deliver antibacterial toxins, or effector proteins, directly into neighbouring competitors, effectively killing or disabling recipient cells^3^. Growing evidence supports a role for the T6SS in determining the success of individual isolates and the composition of the polymicrobial population in a wide variety of host-associated and environmental communities^4, 5, 6^. The T6SS is a dynamic protein nanomachine in which contraction of an intracellular sheath structure anchored on a membrane-bound basal complex propels a cell puncturing device out of the secreting cell and into a neighbouring recipient, or target, cell. This expelled puncturing structure, comprising a tube of Hcp tipped with a spike of VgrG and PAAR proteins, is decorated with multiple effector proteins, which either interact with or are covalently fused to the tube or spike proteins, thereby delivering the effectors into the target cell^7, 8^.

Since the discovery of inter-bacterial competition mediated by the T6SS^9^, an ever-increasing number of T6SS-delivered antibacterial effectors have been reported, with molecular targets that are conserved and essential in bacterial cells^7^. The vast majority of these are enzymatic toxins, including families of effectors which cleave the peptidoglycan cell wall, hydrolyse nucleic acids, degrade membrane phospholipids, deplete NAD or ATP, and modify RNA or proteins by ADP-ribosylation^7, 10, 11^. Self-intoxication and intoxication by incoming effectors delivered by genetically-identical neighbouring cells is prevented through the co-expression of effector-specific immunity proteins. Immunity proteins are localised at the site of action of their cognate effector, bind tightly and specifically to the effector, and, for enzymatic toxins, typically inhibit effector activity by blocking the active site^11^. The mode of action of enzymatic effectors can often be predicted by sequence homology or structural prediction, including, more recently, with the use of AlphaFold2 which can typically generate reliable predictions for soluble enzymatic domains^12^. A small number of T6SS effectors have been reported which, by contrast, act as pore forming toxins in recipient cell membranes, including two shown to form cation-selective pores *in vitro*^13, 14^. However, identification of such effectors has been hampered by the fact that their function is much harder to predict *in silico*, given they typically present as small proteins with no homologues of known structure or function, and we believe that they represent a larger contribution to the pool of antibacterial effectors than is currently appreciated.

*Serratia marcescens* is an opportunistic pathogen which is widespread in diverse environmental niches and represents a significant cause of antibiotic-resistant hospital-acquired infections^15^. The T6SS of the model strain *S. marcescens* Db10 is well characterised and displays potent anti-bacterial and anti-fungal activity. Ten effectors delivered by this T6SS have been identified by secretomic and genetic studies, including anti-bacterial effectors with peptidoglycan amidase (Ssp1 and Ssp2), DNase (Rhs2), and NAD(P)^+^-glycohydrolase (Rhs1) activity, and two anti-fungal effectors^16, 17, 18, 19, 20^. This T6SS also delivers the ion-selective pore forming effector Ssp6, which forms cation-selective pores *in vitro* and, as a consequence, disrupts the inner membrane potential *in vivo*^14^. Another effector, Ssp4, was identified in the T6SS-dependent secretome of *S. marcescens* Db10 and subsequently shown to possess antibacterial activity which was observable upon expression in the periplasm of *E. coli* and neutralised by a cognate immunity protein, Sip4^17^. However, no function could be readily ascribed to Ssp4, given a lack of sequence or structural similarity with any T6SS-associated or other proteins described previously.

Here, we report that Ssp4 forms cation-selective pores in the membrane of susceptible bacterial cells and represents the founding member of a widely-occurring new family of T6SS-dependent pore-forming toxins. Importantly, we show that Ssp4 is active against a wider range of target species than the other cation-selective pore-forming toxin delivered by this T6SS, Ssp6, revealing that not all T6SS antibacterial effectors have broad spectrum activity. We further demonstrate that the ion selectivities of the pores formed by Ssp4 and Ssp6 are distinct and provide the first high resolution model of a membrane pore formed by a T6SS-delivered effector. Our data support a model whereby possession of two distinct T6SS-dependent pore-forming toxins ensures effective de-energisation of both closely- and distantly-related competitors.

## Results

### The properties of Ssp4 and Sip4 are consistent with Ssp4 being a membrane-targeting effector

Ssp4 is encoded with its cognate immunity protein, Sip4, within a three gene insertion in the isoleucine, leucine and valine (*ilv*) gene cluster of *S. marcescens* Db10 (Fig. 1a). AlphaFold2^12^ was used to generate a high confidence structural prediction for Ssp4, comprising 13 α-helices and one β-sheet (pLDDT value 85.9, Fig. 1b,c). Regions within three of these α-helices were predicted to represent transmembrane helices by MEMSAT (Fig. 1b,c), indicating that Ssp4 may have the ability to integrate into membranes. However, no Ssp4 homologues of known or predicted function were identified in sequence databases and no convincing structural homologues were retrieved when the predicted structure of Ssp4 was used to search the Protein Data Bank.

**Figure 1.**
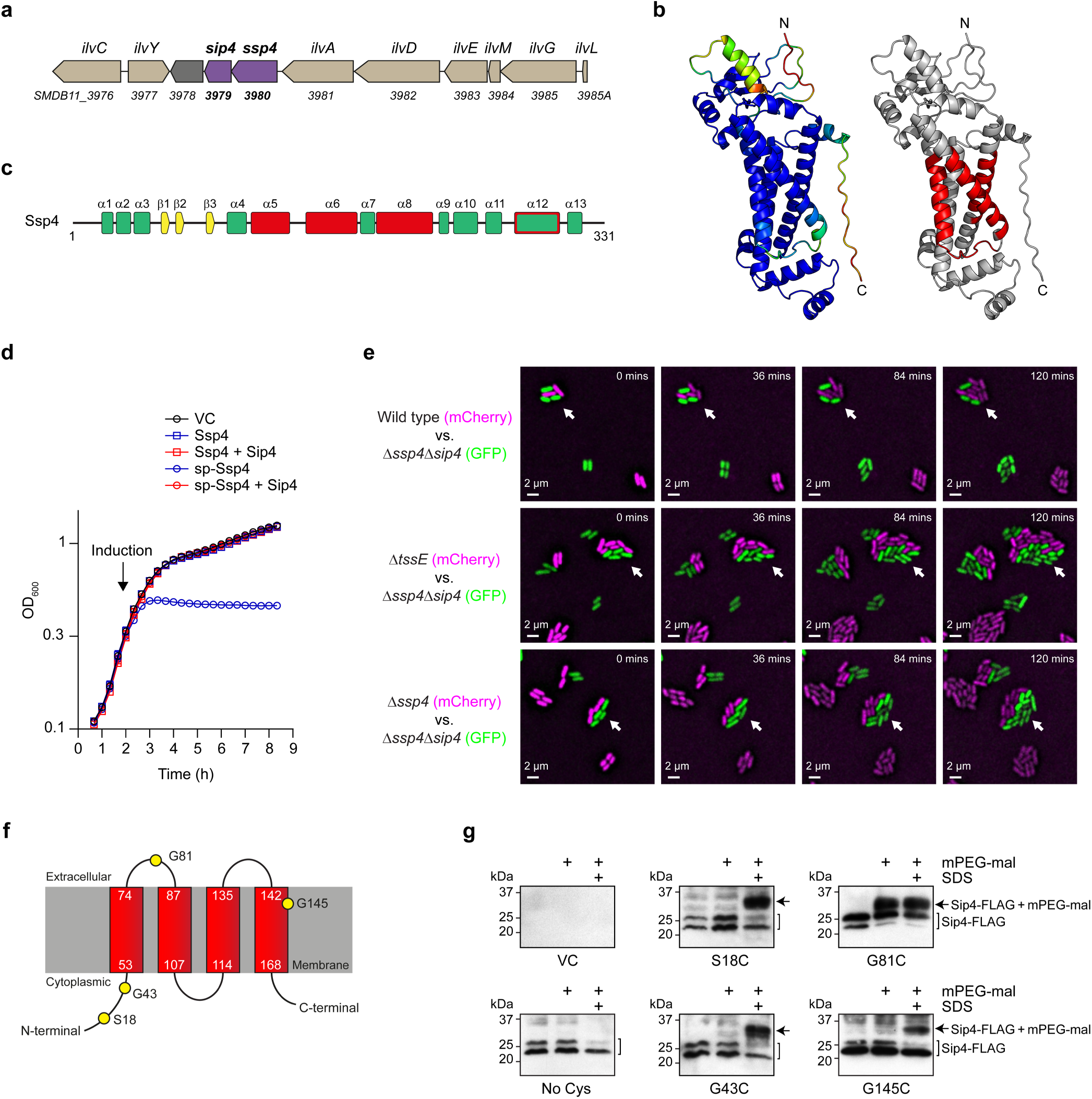
Ssp4 is a non-lytic antibacterial toxin that acts from within the periplasm and is neutralised by the integral inner membrane immunity protein Sip4. (a) Genomic context of the genes encoding Ssp4 and Sip4 in *S. marcescens* Db10, with genomic identifiers (SMDB11_xxxx) below each gene and gene names above (*ilv*, isoleucine, leucine, and valine genes). (b) Structure of Ssp4 predicted by AlphaFold2. Left, structure is coloured by pLDDT value, spectrum from red (<50) to blue (>90). Right, regions predicted to form transmembrane helices by MEMSAT2 are highlighted in red. (c) Secondary structure elements in the predicted structure of Ssp4, with α-helical regions predicted to include transmembrane helices by MEMSAT coloured red; the predicted structure of Ssp4 indicates that helix α12 should also cross the membrane as shown by a red outline. (d) Growth of *E. coli* MG1655 carrying the vector control (VC, pBAD18-Kn) or plasmids directing the expression of Ssp4 or Ssp4 fused with an N-terminal OmpA signal peptide (sp-Ssp4), either alone or with Sip4. Gene expression was induced by addition of 0.2% L-arabinose at the time indicated. Points show mean +/- SEM (n=3 biological replicates). (e) Ssp4-mediated growth inhibition observed by time-lapse fluorescence microscopy. An Ssp4-susceptible target, *S. marcescens* Δ*ssp4*Δ*sip4* expressing cytoplasmic GFP (green), was co-cultured with wild type (WT) or mutant (Δ*tssE* or Δ*ssp4*) attacker strains expressing cytoplasmic mCherry (magenta) for 2 h. Arrows highlight example microcolonies where attacker and target cells are in contact. Scale bar 2 µm. Images are representative of three independent experiments including at least nine frames per attacker strain. (f) Predicted membrane topology of Sip4 generated by MEMSAT with amino acids selected for cysteine substitution represented as yellow circles. (g) Cells of *S. marcescens* Δ*ssp4*Δ*sip4* carrying the vector control (VC, pSUPROM) or plasmids directing the expression of Sip4-FLAG with native Cys residues mutated to Ala (No Cys) or derivatives carrying the Cys substitutions indicated, were treated with mPEG-MAL in the presence or absence of SDS. Sip4-FLAG and Sip4-FLAG-mPEG-MAL species were detected by immunoblotting.

In order to investigate the mode of toxicity of Ssp4, *E. coli* MG1655 was transformed with plasmids directing the expression of Ssp4, either retained in the cytoplasm or fused to an N-terminal signal peptide for export to the periplasm (sp-Ssp4). Inducing expression of sp-Ssp4 inhibited further growth but did not cause a drop in optical density, suggesting that Ssp4 does not cause lysis of intoxicated cells (Fig. 1d). No inhibition was observed when Ssp4 was retained in the cytoplasm or when Sip4 was co-expressed. Next, we determined whether the action of Ssp4 results in growth inhibition at a single cell level when it is delivered into neighbouring cells by the T6SS. Time-lapse fluorescence microscopy was used to observe co-cultures between cells of wild type, T6SS-inactive (Δ*tssE*), or Δ*ssp4* strains of *S. marcescens* Db10 expressing mCherry (‘attacker’) and an Ssp4-susceptible ‘target’ strain, Db10 Δ*ssp4*Δ*sip4* expressing GFP (this mutant lacks the protection normally conferred by Sip4). Target cells in contact with wild type attacker cells generally failed to proliferate and divide, whilst isolated target cells or target cells in contact with T6SS-deficient or Ssp4-lacking attackers proliferated indistinguishably from attacker cells (Fig. 1e). Given also that no target cell lysis events were observed, our combined data are consistent with Ssp4 intoxication causing growth inhibition but not cell lysis.

The subcellular localisation of T6SS immunity proteins typically indicates the site of action of the cognate effector^3^. Using MEMSAT, Sip4 was predicted to contain four transmembrane helices, with an N- and C-terminal ‘in’ topology (Fig. 1f). This prediction was consistent with a low confidence AlphaFold model in which Sip4 folds into a bundle of seven α-helices (pLDDT 64.9; Suppl. Fig. 1) and suggested that Sip4 is an integral membrane protein. To confirm the membrane insertion and topology of Sip4, fully functional variants of Sip4 with no cysteine residues or selected single cysteine substitutions were expressed in the Db10 Δ*ssp4*Δ*sip4* mutant (Fig. 1f, Suppl. Fig. 2) and cells were incubated with the membrane impermeant, Cys-reactive reagent mPEG-malemide (mPEG-Mal). Sip4 variants with cysteine substitutions in residues predicted to be located in the cytoplasm (S18C, G43C) or a transmembrane helix (G145C) were only labelled with mPEG-Mal in the presence of SDS, confirming that these amino acid positions are not exposed extra-cytoplasmically and only become available for labelling upon inner membrane disruption (Fig. 1g). In contrast, the G81C variant could be labelled in the absence of SDS, confirming that Gly 81 is exposed in the periplasm (Fig. 1g). These data confirm that Sip4 is an integral inner membrane protein and are consistent with the predicted topology. Taken together, the predicted α-helical structure of Ssp4, its non-lytic and periplasmic toxicity, and the integral membrane location of its immunity protein, suggested that Ssp4, like Ssp6, was likely to be a membrane-targeting toxin.

### Ssp4 and Ssp6 display distinct target specificities during T6SS-mediated competition

Next we compared the ability of Ssp4 and the previously-characterised membrane-targeting effector Ssp6 to intoxicate different bacterial species upon delivery by the T6SS. Initially, we compared the T6SS-dependent anti-bacterial activity of Δ*ssp4* and Δ*ssp6* mutants with that of wild type *S. marcescens* Db10 against *Pseudomonas fluorescens* and *E. coli*, by determining the recovery of viable target cells following co-culture with the attacker strain of *S. marcescens*. While the Δ*ssp4* mutant showed a small reduction in activity against *P. fluorescens*, in the other cases no significant difference with wild type Db10 was observed (Fig. 2a,b). This is not unexpected, given that there are at least seven other anti-bacterial effectors delivered by the Δ*ssp4* and Δ*ssp6* mutants. Therefore, to observe the impact of Ssp4 and Ssp6 in isolation, we separately re-introduced the gene for each effector into *S. marcescens* Db10 lacking all known anti-bacterial effectors (strain Δ9). Against *E. coli*, both Δ9 + *ssp4* and Δ9 + *ssp6* displayed considerable anti-bacterial activity compared with Δ9, showing that both effectors can act against this species and that Ssp4 is the more potent (Fig. 2a). Ssp4 also showed substantial activity in *S. marcescens* when Db10 or Δ9 + *ssp4* were co-cultured with the Δ*ssp4*Δ*sip4* mutant (Fig. 2c). Unexpectedly, whilst Ssp4 also effectively intoxicated *P. fluorescens*, Ssp6 showed no detectable activity against this target species (Fig. 2b), revealing that the two effectors have an overlapping but distinct range of target species.

**Figure 2.**
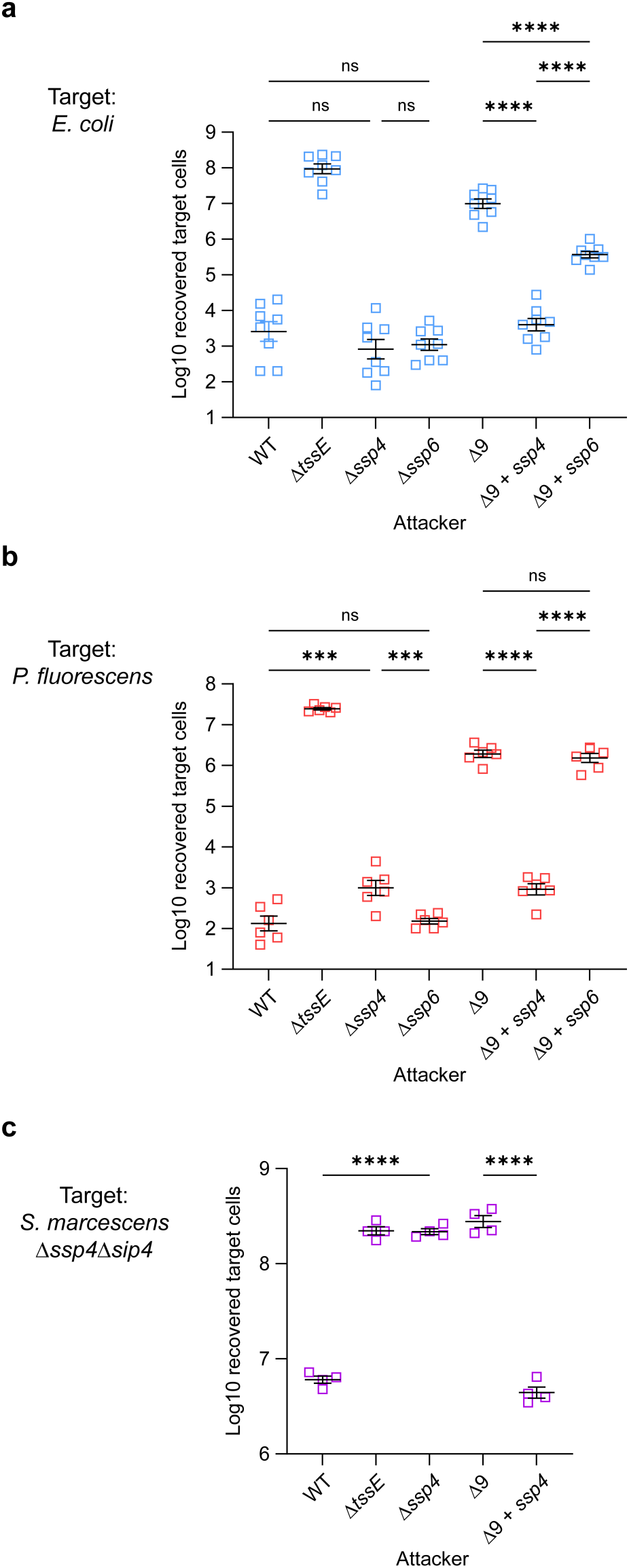
Ssp4 and Ssp6 differ in their activity against different target species when delivered by the T6SS. T6SS-dependent anti-bacterial activity of wild type (WT) or mutant (Δ*tssE*, Δ*ssp4*, Δ*ssp6*, Δ9, Δ9 + *ssp4* and Δ9 + *ssp6*) strains of *S. marcescens* Db10, as indicated, against (a) *E. coli* BW25112, (b) *P. fluorescens* 55 or (c) *S. marcescens* Db10 Δ*ssp4*Δ*sip4* target strains. The Δ9 mutant lacks all known anti-bacterial effectors in Db10; Δ9 + *ssp4* and Δ9 + *ssp6* have the respective effector reintroduced. Recovery of target cells was enumerated following co-culture of attacker and target at an initial ratio of 1:1 for 4 h. Data are presented as mean +/- SEM with individual data points overlaid (n=8, n=6 and n=4 biological replicates for panels a, b and c, respectively; **** p<0.0001, *** p<0.001, ns not significant, one-way ANOVA with Tukey’s test; for clarity, only selected comparisons are displayed).

### Ssp4 causes loss of membrane potential in susceptible cells

Given the phenotypic similarity of Ssp4 with other membrane-targeting effectors, we investigated whether Ssp4 affects the membrane potential or permeability of intoxicated cells. We co-cultured attacker strains delivering Ssp4 (wild type Db10 or Δ9 + *ssp4*) with the Ssp4-susceptible target Δ*ssp4*Δ*sip4*, and stained the total mixed population with the voltage-sensitive dye DiBAC_4_(3) and propidium iodide (PI). Around 15-20% of the total population were positive for DiBAC_4_(3) fluorescence, indicating that the cells had become depolarised. However these cells did not simultaneously stain with PI, indicating that there was no permeabilisation or loss of membrane integrity (Fig. 3a). The population of DiBAC_4_(3)-positive, depolarised cells was not observed when the attacking cells were unable to deliver Ssp4 (Δ*tssE*, Δ*ssp4* and Δ9 mutants), confirming that it represents Ssp4-intoxicated cells. Similarly, heterologous expression of sp-Ssp4 in *P. fluorescens* induced membrane depolarisation but not permeabilisation (Fig. 3b). Therefore Ssp4, like Ssp6^14^, disrupts the inner membrane potential in intoxicated cells, without the formation of large unspecific pores or loss of bilayer integrity. However, no DiBAC_4_(3) or PI staining above control was observed on expression of sp-Ssp6 in *P. fluorescens* (Fig. 3b), consistent with the lack of Ssp6 activity observed against *P. fluorescens* in co-culture (Fig. 2b). When cells of *P. fluorescens* expressing sp-Ssp4 or sp-Ssp6 were fractionated, Ssp4 could be observed within the total membrane fraction, confirming that Ssp4 interacts directly with target cell membranes (Fig. 3c). At least small amounts of Ssp6 could also be observed associated with the total membrane fraction in this system.

**Figure 3.**
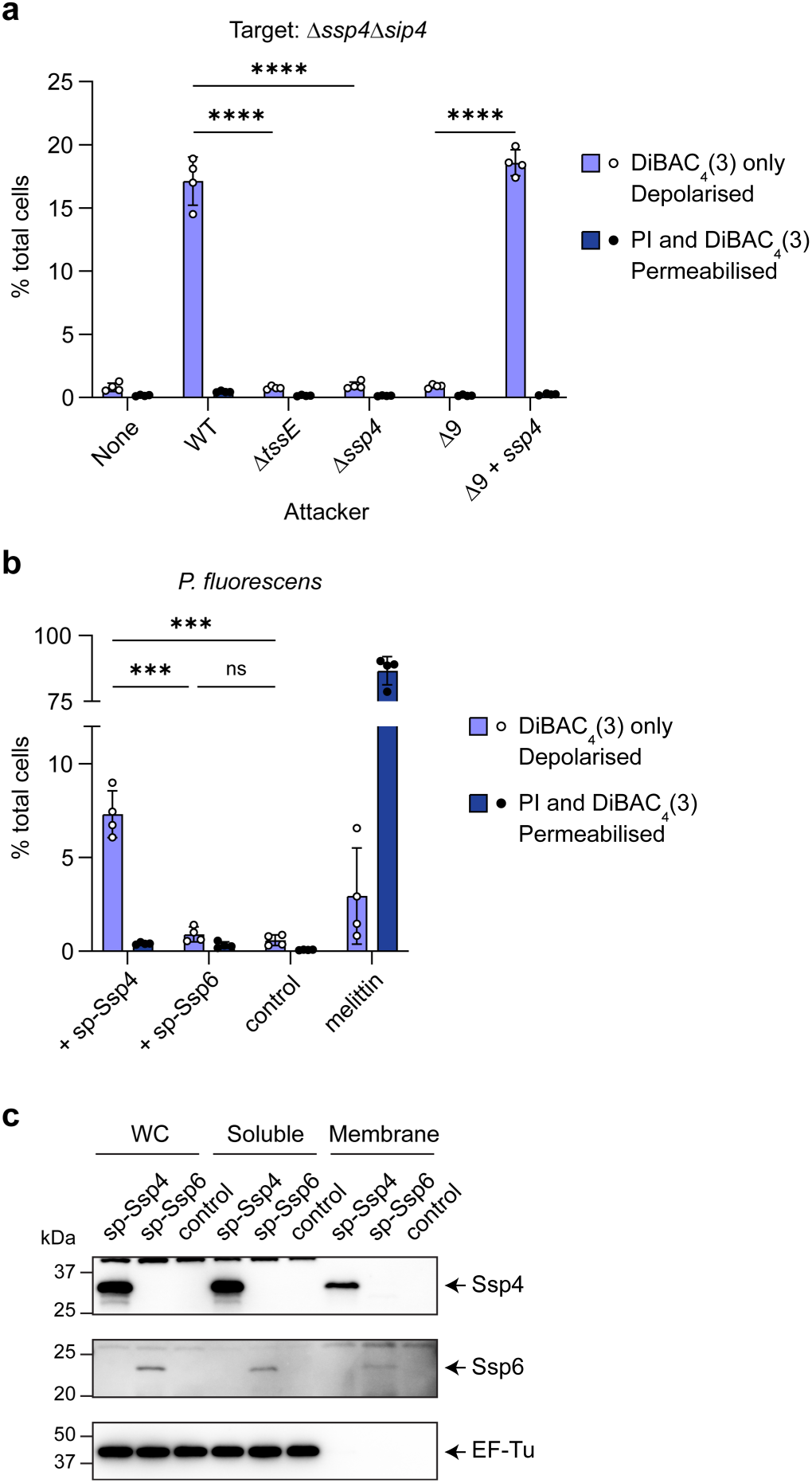
Intoxication by Ssp4 causes loss of membrane potential. (a) The Ssp4-susceptible mutant of *S. marcescens* Db10, Δ*ssp4*Δ*sip4,* was co-cultured with wild type (WT) or mutant (Δ*tssE,* Δ*sip4*, Δ9 and Δ9 + *ssp4*) attacker strains, as indicated, at a starting ratio of 1:2 for 4 h, then the mixed population was stained with DiBAC_4_(3) and propidium iodide (PI) and analysed by flow cytometry to determine membrane potential and permeability, respectively. The percentage of cells in the total bacterial population identified as being depolarised (positive for DiBAC_4_(3) staining, negative for PI staining), or permeabilised (positive for both DiBAC_4_(3) and PI staining) were quantified. (b) Cells of *P. fluorescens* carrying a chromosomal insertion directing the expression of Ssp4 or Ssp6 fused with an N-terminal signal peptide (sp-Ssp4 or sp-Ssp6) under the control of P_Rha_, or P_Rha_ alone (control) were induced by the addition of 0.05% rhamnose and the percentage of cells depolarised or permeabilised was determined as in panel a. Cells treated with melittin acted as a positive control for permeabilisation. For panels a and b, data are presented as mean ± SEM with individual data points overlaid (n=4 biological replicates; **** p<0.0001, *** p<0.001, ns not significant, one-way ANOVA with Tukey’s test; for clarity, only selected comparisons are displayed). (c) Immunoblot detection of Ssp4, Ssp6 and EF-Tu (cytoplasmic control) in whole cell (WC), soluble or total membrane fractions of *P. fluorescens* expressing sp-Ssp4 or sp-Ssp6 as in panel b.

### Ssp4 forms ion-selective membrane pores *in vitro*

The observation that Ssp4 intoxication causes inner membrane depolarisation suggested that this protein, like Ssp6^14^, may form ion-selective pores, leading to disruption of the membrane potential through ion leakage. To test the ability of Ssp4 to form membrane pores, purified monomeric Ssp4 protein (Suppl. Fig. 4) was incorporated into artificial lipid bilayers under voltage-clamp conditions. In symmetrical KCl (510 mM KCl in *cis* and *trans* chambers), Ssp4 generated a measurable current when the membrane was clamped at voltages of > 40 mV or < -40 mV, confirming that Ssp4 is capable of ion permeation (Fig. 4a,b). Between 20 mV and -20 mV, the signal-to-noise ratio was too high to accurately determine current measurements, although pore openings remained visible (Fig. 4a). In symmetrical 510 mM KCl solutions, Ssp4 displayed a single channel conductance of 18.4 ± 0.64 pS (mean ± SD; n = 4), consistent with the conduction of ions through the Ssp4 pore. In order to investigate whether Ssp4 is permeable to cations, anions or both, a current/voltage (*I*/*V*) relationship in non-symmetrical conditions was determined and used to calculate the reversal potential (*E_rev_*) of the Ssp4 pore. Under these conditions, the reversal potential was -14.8 ± 1.6 mV (mean ± SD; n = 4), a value that is closer to the predicted equilibrium potential (calculated using the Nernst equation) of K^+^ (−22.8 mV) than Cl^-^ (+22.8 mV), indicating a preference for cations over anions (Fig. 4b).

**Figure 4.**
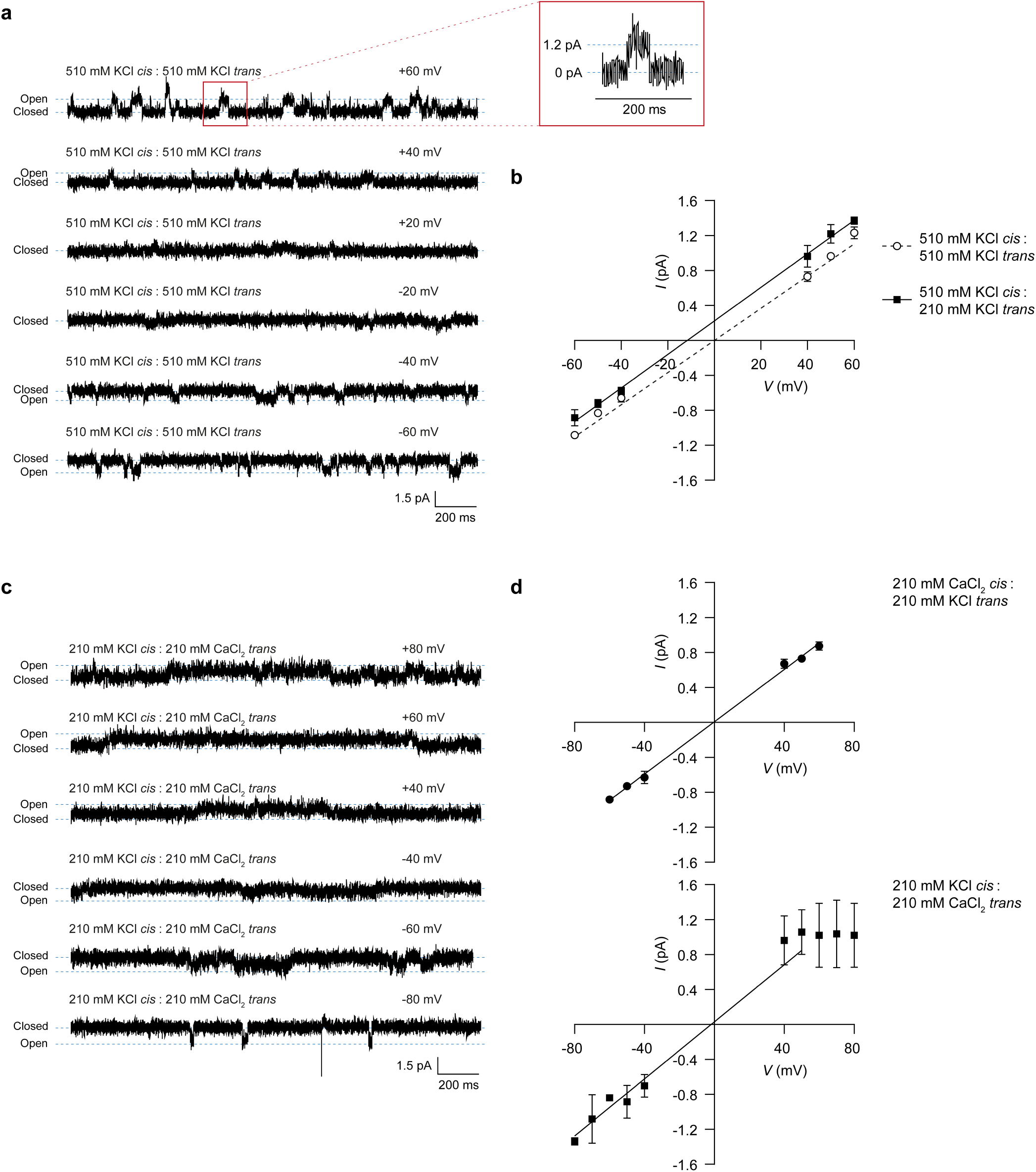
Ssp4 forms ion-selective pores in planar lipid bilayers *in vitro*. Purified Ssp4 was incorporated into artificial lipid bilayers under voltage-clamp conditions and the resulting current measured. (a) Representative single-channel experiment in which Ssp4 was added to the *cis* chamber with 510 mM KCl as the source of permeant ions. (b) Current-voltage relationship for Ssp4 with KCl as the source of permeant ions. Solid line, symmetrical 510 mM KCl; dotted line, 510 mM KCl *cis* chamber, 210 mM KCl *trans* chamber. (c) Representative single-channel experiment in which Ssp4 was added to the *cis* chamber with 210 mM KCl *cis* and 250 mM CaCl_2_ *trans* as the source of permeant ions. (d) Current-voltage relationship with KCl and CaCl_2_ as the source of permeant ions. Top, 210 mM CaCl_2_ *cis* chamber, 210 mM KCl *trans* chamber; bottom, 210 mM KCl *cis* chamber, 210 mM CaCl_2_ *trans* chamber. Data are displayed as mean ±SD from 3 or 4 independent experiments.

In order to establish whether the Ssp4 pore has higher selectivity for monovalent or divalent cations, the relative permeability of K^+^ and Ca^2+^ was examined. When Ca^2+^ was the permeant ion in the *cis* chamber and K^+^ the permeant ion in the *trans* chamber, the reversal potential was close to zero (-1.44 ± 1.7 mV; mean ± SD; n = 3), implying that Ssp4 transports monovalent and divalent cations equally well (Fig. 4d). When CaCl_2_ was in the *trans* chamber and KCl in the *cis* chamber, rectification was apparent at voltages > 50 mV, but not at negative potentials, indicative of a change in the behaviour of the pore (Fig. 4c,d). Additionally, rectification of the channel by high Ca^2+^ only in the *trans* chamber suggests that Ssp4 incorporates into the membrane in a fixed orientation. The reversal potential of the pore was calculated through point-to-point analysis of the linear current-voltage relationship at -50 mV and + 50 mV, giving a value of -3.1 ± 4.6 (mean ± SD; n = 3). The single channel conductance of the pore also changed when Ca^2+^ was present in the *trans* chamber, increasing from 15.0 ± 0.40 pS (mean ± SD; Ca^2+^ ions in *cis*) to 19.4 ± 3.2 pS (mean ± SD; Ca^2+^ ions in *trans*) suggesting that the channel may undergo a Ca^2+^-dependent conformational change.

### Structural modelling and molecular dynamics simulation suggest that Ssp4 forms tetrameric pores in membranes

In order to investigate the potential structure of the pore formed by Ssp4 in target cell membranes, structural predictions for Ssp4 homo-multimers containing two, three, four, six or eight Ssp4 monomers were generated using AlphaFold2^12^, based on full length Ssp4 (Ssp4_1-331_) or a truncated version lacking regions disordered in the predicted structure of the monomer (Ssp4_114-331_). The ability of the resulting structural models to insert into a POPE/POPG lipid bilayer was predicted using MemProtMD^21^. Following manual examination of the subset of models predicted to form a transmembrane complex, a tetrameric assembly was selected as the most likely structure for an ion-selective pore and a final model based on the predicted tetrameric structure of Ssp4_114-331_ with manual removal of the final 29 amino acids (Ssp4_114-302_) was taken forward for analysis by molecular dynamics simulation. In this model, the four Ssp4_114-302_ proteins retain a similar structure to that predicted for the corresponding regions of the Ssp4 monomer in isolation (Fig. 1b) and form a ring-like assembly with a central channel and four transmembrane helices contributed by each Ssp4 (Fig. 5a-c). Based on MEMSAT predictions and the fact that Ssp4 is active from the periplasm (Fig. 1d), which suggests that Ssp4 enters the membrane from the periplasmic side, we predict that the pore is oriented with its N- and C-termini (including the ∼113 amino acid N-terminal domain truncated from the model) in the periplasm.

**Figure 5.**
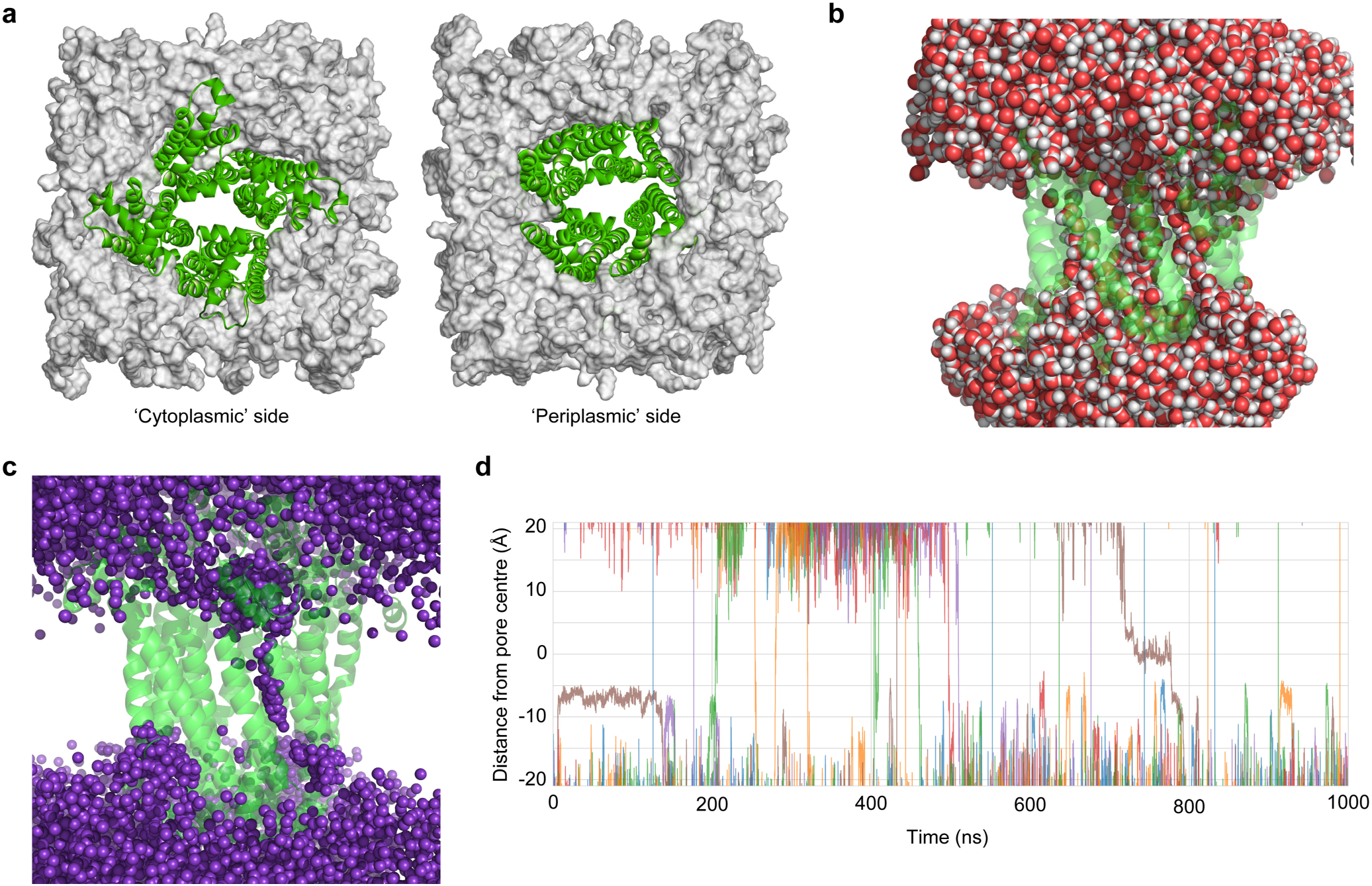
Molecular dynamics simulation of ion permeation through a tetrameric Ssp4 pore. (a) Model structure of the Ssp4_114-302_ tetramer in a POPC lipid bilayer. Left: view of the pore from the side predicted to be on the cytoplasmic side of the membrane *in vivo*, including helices α10 and α11 which are peripheral to the membrane; right: view of the pore from the side on which the N- and C-termini of Ssp4 are located, predicted to be on the periplasmic side of the membrane *in vivo*. (b) Hydration of the Ssp4_114-302_ tetramer in the membrane. (c) An overlay of K^+^ ion positions from a 1 µs simulation reveals a preferred ion permeation pathway across the pore formed by Ssp4. (d) Permeation traces of K^+^ ions along the membrane axis.

To investigate the structural stability of the Ssp4 tetrameric model and its permeability for ions, we performed atomistic molecular dynamics simulations of membrane-embedded, tetrameric Ssp4_114-302_. Our initial set of molecular dynamics simulations of the Ssp4_114-302_ tetramer embedded in a model lipid bilayer (Fig. 5a) showed structural stability with a root-mean-square deviation between 2.5-4 Å over a simulated time of 300 ns (Suppl. Fig. 5). To investigate the model’s permeability for ions, we next performed simulations of Ssp4 at a membrane voltage of 100 mV. Each simulation had a length of 1 µs and was three-fold replicated. Across all simulations taken together, we observed the permeation of 29 K^+^ ions and 4 Cl^-^ ions, confirming both the ability of the membrane-inserted Ssp4 tetramer to conduct ions and its cation selectivity. The conductance calculated from these simulations was 18 ± 6 pS, in good agreement with the experimental conductance of 18.4 pS observed above. Figure 5b shows the typical hydration pattern of the Ssp4_114-302_ tetramer inserted in the membrane, revealing multiple narrow water channels. These pores allow ions to traverse the membrane under voltage (Fig. 5c,). Although we cannot rule out effects related to the truncation of the Ssp4 sequence, the central water channel remains hydrated throughout all our simulations, highlighting the hydrophilic character of the interior of the tetrameric pore.

### Ssp4 intoxication is accompanied by increased levels of intracellular reactive oxygen species

Given that Ssp4 and Ssp6 are both ion-selective pore-forming effectors, we next asked if they cause the same response in intoxicated cells. Many antibiotics and other stressors, including phage infection and T6SS attack, have been reported to cause production of reactive oxygen species (ROS) in bacterial cells^22, 23^ ^24, 25^. To determine if ROS generation occurs following Ssp4 intoxication, sp-Ssp4 was expressed in *E. coli* and ROS levels quantified by OxyBURST Green staining. A significant increase in staining was recorded 2 and 3 h after induction of sp-Ssp4, indicating that the cells had entered a state of oxidative stress, but this was prevented by co-expression of Sip4 (Fig. 6a,b). In contrast, no specific increase in OxyBURST staining was observed when *E. coli* was intoxicated with sp-Ssp6 or the cytoplasmic-acting effector Ssp5 (Fig. 6c,d), suggesting that the generation of ROS is a specific consequence of Ssp4 intoxication.

**Figure 6.**
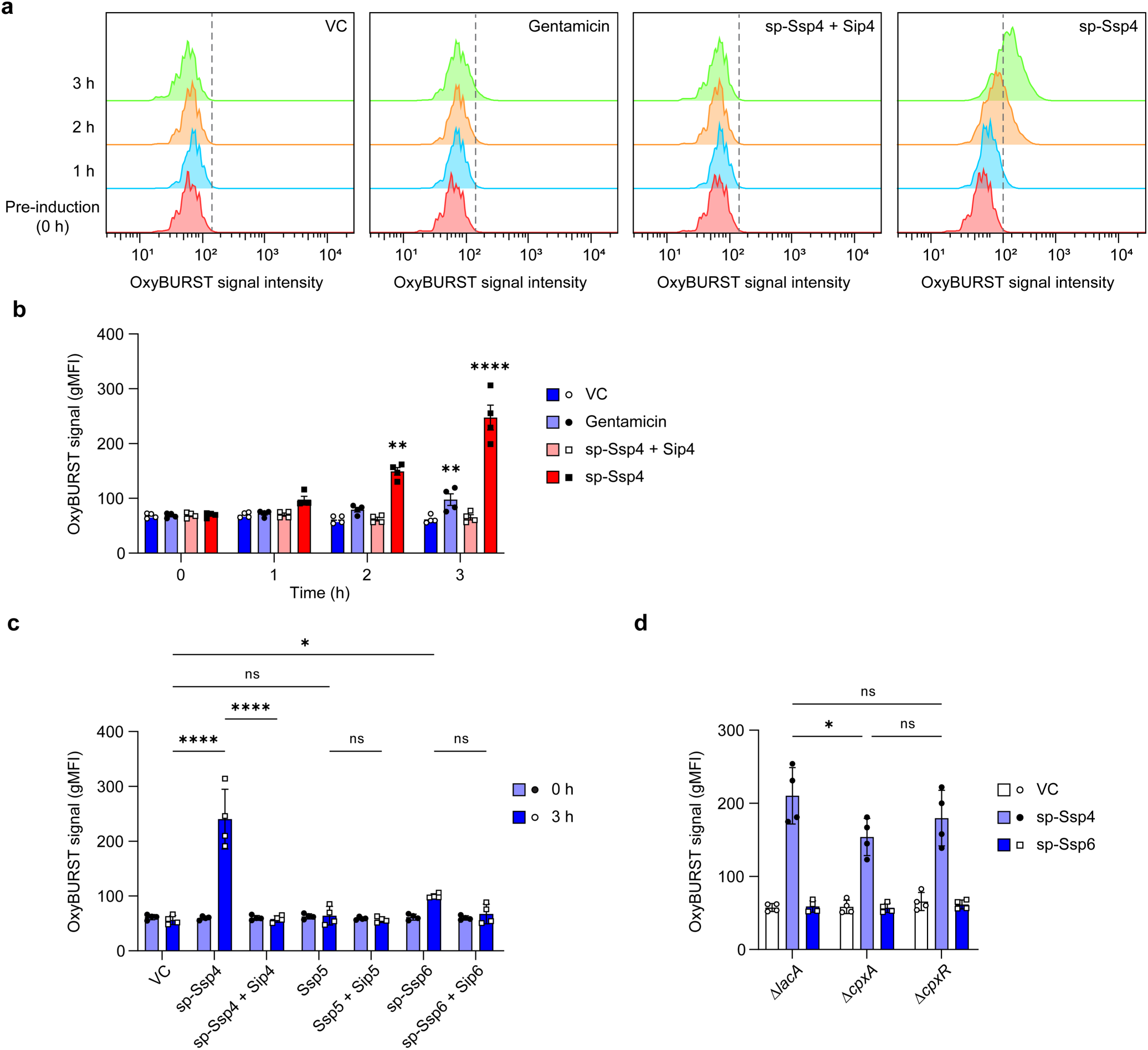
Ssp4 intoxication is accompanied by an increase in the level of intracellular reactive oxygen species (ROS). (a) Representative flow cytometry histograms showing the OxyBURST Green signal intensity over time from cells of *E. coli* MG1655 carrying the vector control (VC, pBAD18-Kn) or plasmids directing the expression of Ssp4 fused with an N-terminal signal peptide (sp-Ssp4), either alone or with Sip4, following induction with 0.2% L-arabinose. Cells were separately exposed to 5 µg/ml gentamicin for comparison. (b) Quantification of OxyBURST Green signal from the experiment in panel a, with data presented as mean ± SEM, with individual data points overlaid (n=4 biological replicates; **** p<0.0001, ** p<0.01, compared with pre-induction levels; repeated measures ANOVA with Dunnett test. gMFI, geometric mean fluorescence intensity. (c, d) Quantification of OxyBURST Green intensity from (c) *E. coli* MG1655 carrying plasmids directing expression of sp-Ssp4, Ssp5 and sp-Ssp6, with or without their cognate immunity proteins, pre-induction and following 3 h induction; or (d) strains of *E. coli* BW25113 (Δ*lacA*, Δ*cpxA*, Δ*cpxR*) carrying plasmids directing expression of sp- Ssp4 or sp-Ssp6, following 3 h induction. Data are presented as mean +/- SEM with individual data points overlaid (n=4 biological replicates; **** p<0.0001, * p<0.05, ns not significant, one-way ANOVA with Tukey’s test; for clarity, only selected comparisons are displayed).

The generation of ROS can be triggered via number of different pathways and several gene deletion mutants have been reported to show reduced susceptibility to antibiotics or toxins as a result of reduced ROS production ^25, 26, 27, 28^. As Ssp4 targets the membrane, we investigated the potential contribution of the CpxAR-dependent envelope stress response which is triggered in response to membrane disruption and has been implicated in ROS-dependent toxicity^26, 27, 28^. The sp-Ssp4 protein was expressed in *E. coli* BW25113 Δ*cpxA* and Δ*cpxR* mutants, as well as in a Δ*lacA* control mutant derived from the same collection^29^. The Δ*cpxA* mutant showed a modest decrease in OxyBURST signal intensity upon Ssp4 intoxication, indicating that CpxA activation partly contributes to Ssp4-induced ROS generation (Fig. 6d). However this effect was not sufficient to provide the Δ*cpxA* mutant with measurable resistance against Ssp4 in the context of T6SS-mediated competition (Suppl. Fig. 6), while the Δ*cpxR* mutant showed similar resistance towards Ssp4 as the Δ*lacA* control in both assays.

### Use of Tn-seq to identify mutants resistant to individual effectors reveals that MucA disruption protects *P. fluorescens* against T6SS attack

We next aimed to identify mutations conferring resistance to Ssp4 intoxication in an unbiased manner using transposon insertion site sequencing (Tn-seq) (Fig. 7a). A transposon insertion library of *P. fluorescens* was co-cultured with *S. marcescens* Db10 Δ9 or Δ9 + *ssp4* and the recovered *P. fluorescens* population was subjected to Tn-seq in order to identify genes whose inactivation led to an over-representation of the corresponding mutants in the *P. fluorescens* population exposed to Ssp4 delivery compared with that exposed to the Δ9 control. The Δ9 + *ssp6* strain was also included and the equivalent comparison with Δ9 performed, as an example of a pore-forming effector which does not detectably harm *P. fluorescens*. The Tn-seq experiment revealed one gene whose disruption resulted in increased relative survival under conditions of Ssp4 intoxication, *moaC*, and two genes whose disruption resulted in increased relative survival under Ssp6 intoxication, *xcpW* and *mucA* (FDR <0.05, Fig. 7b, Table 1, Suppl. Dataset 1). MoaC is required for molybdenum cofactor biosynthesis^30^, XcpW is a minor pseudopilin in the Type II secretion system^31^, and MucA is a negative regulator of the sigma factor AlgU, preventing activation of the AlgU regulon in response to cell envelope stress^32^. A limited number of genes whose disruption resulted in decreased relative survival under Ssp4 intoxication conditions were identified (Table 1); two of these, *01041* and *00266*, encoding proteins of unknown function, were selected for further study, along with *moaC*, *xcpW* and *mucA*. Transposon insertions were observed across the full length of *moaC*, *xcpW*, *01041* and *00266* (Suppl. Fig. 7). However, the first 200 bp of *mucA* was devoid of transposon insertions, suggesting that this region of the gene is required for bacterial viability, consistent with the observation that the N-terminal AlgU binding domain of MucA is essential for viability in *P. aeruginosa*^32^.

**Figure 7.**
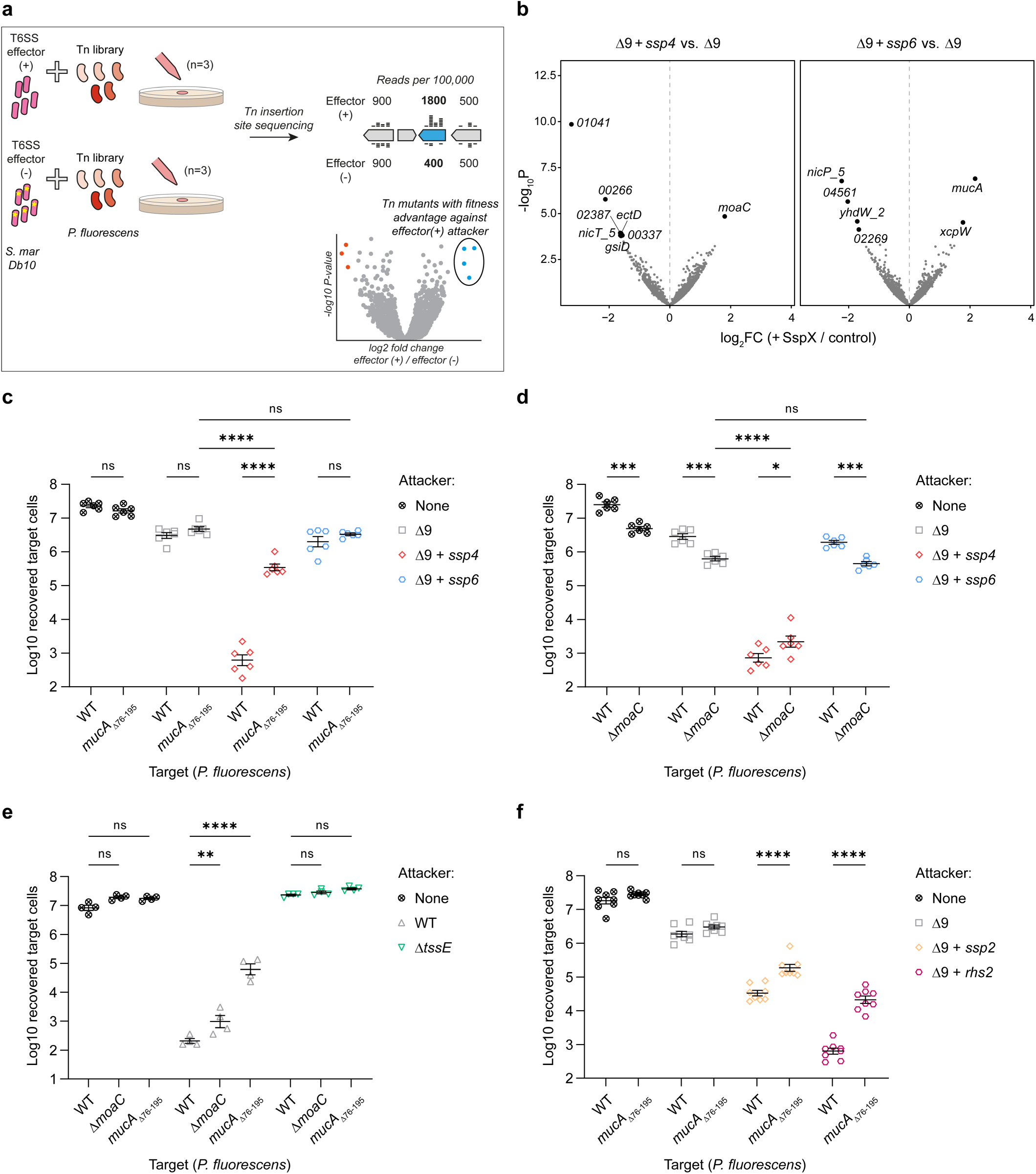
Use of Tn-seq to identify mutants resistant to Ssp4 or Ssp6 reveals that MucA disruption protects *P. fluorescens* against the T6SS. (a) Schematic illustration of the Tn-seq experiment. *S. marcescens* Db10 lacking known anti-bacterial effectors (effector (-), strain Δ9) or delivering only one effector (effector (+), strains Δ9 + *ssp4* or Δ9 + *ssp6*) was co-cultured with a saturated transposon insertion library of *P. fluorescens* 55 for 4 h. Genomic DNA from the recovered total population was subjected to Illumina sequencing using transposon specific primers and the number of reads mapping to each *P. fluorescens* gene determined, indicating the relative abundance of *P. fluorescens* mutants with insertions in each gene within the final population. Comparing the relative abundance of mutants with Tn insertions in particular genes between co-culture with effector (+) vs. effector (-) attackers, by determining the fold change in normalised Tn insertion frequency between effector (+) and (-) for each gene, those genes whose disruption results in a fitness advantage against T6SS-mediated intoxication by the effector can be identified (blue points). (b) Volcano plots summarising the change in recovery of transposon insertion mutants between control (Δ9) and Ssp4-delivering (left) or Ssp6-delivering (right) attackers, on a per gene basis. Log2 fold change in normalized read count is plotted against -log10 p-value and genes significantly altered between condition are highlighted (FDR <0.05, n=3 biological replicates). (c-f) Recovery of wild type (WT) or defined mutants (*mucA*_Δ76-195_ or Δ*moaC*) of *P. fluorescens* following co-culture with attacking strains of *S. marcescens* Db10 as indicated (where Δ9 + *ssp2* and Δ9 + *rhs2* intoxicate using only Ssp2 or Rhs2, respectively). None, no attacker. Data are presented as mean ± SEM with individual data points overlaid (n=6, 4 or 8 biological replicates for panels c-d, e, and f, respectively; **** P<0.0001, *** P<0.001, * P<0.05, ns, not significant; one-way ANOVA with Tukey’s test; for clarity, only selected comparisons are displayed). Panels c and d form part of the larger experiment depicted in Supplementary Figure 8.

**Table 1.**
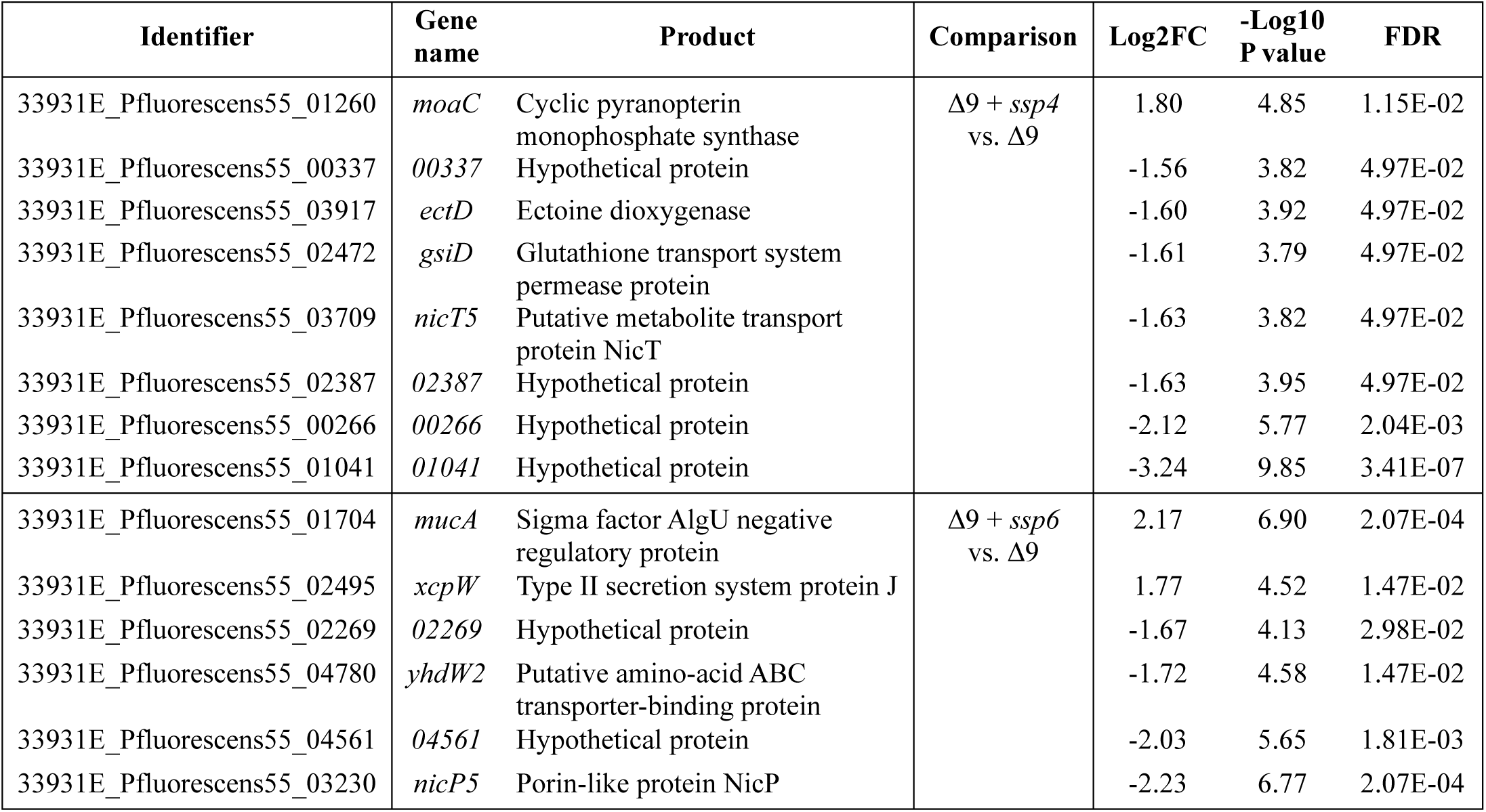
Genes of *P. fluorescens* 55 in which mutants carrying transposon insertions were significantly enriched or depleted in the presence of *S. marcescens* delivering Ssp4 or Ssp6. Genes are included if the number of sequencing reads derived from transposon insertions within the gene was significantly altered between the attacking strains compared, based on an FDR value <0.05. Table summarises data from three biologically-independent co-cultures between a *P. fluorescens* 55 transposon insertion library and *S. marcescens* strains Δ9 (lacking known anti-bacterial effectors), Δ9 + *ssp4* (delivering Ssp4) or Δ9 + *ssp6* (delivering Ssp6). FC, fold change; FDR, false discovery rate.

To validate the results of the Tn-seq experiment, mutants with in-frame deletions of *moaC*, *xcpW*, *01041*, *00266* and the region encoding the C-terminal 120 amino acids of MucA (*mucA*_Δ76-195_) were assessed for susceptibility to Ssp4 and Ssp6 in the standard co-culture assay. The *mucA*_Δ76-195_ mutant displayed a 2-3 log_10_ increase in survival compared with wild type *P. fluorescens* against Δ9 + *ssp4* (Fig. 7c). This finding prompted us to re-examine the TnSeq data and, indeed, *mucA* was the third most increased gene in terms of number of insertions in Δ9 + *ssp4* vs Δ9 (2.7x, p <0.001) but missed the FDR <0.05 cut-off in this case (Suppl. Dataset 1, Suppl. Fig. 7). Recovery of the Δ*moaC* mutant was significantly lower than the wild type under all conditions except when exposed to Δ9 + *ssp4*, indicating this mutation may confer a slight fitness advantage under conditions of Ssp4 intoxication (Fig. 7d). The other *P. fluorescens* mutants showed very small or no differences in resistance to Ssp4 and Ssp6 intoxication (Suppl. Fig. 8).

To determine if the increased resistance of *mucA*_Δ76-195_ was specific to intoxication by Ssp4, this mutant was competed against wild type *S. marcescens* Db10, with a full set of T6SS effectors, and strains delivering unrelated individual effectors, namely the peptidoglycan amidase Ssp2 (Δ9 + *ssp2*) and the DNase Rhs2 (Δ9 + *rhs2*). The *mucA*_Δ76-195_ mutant was considerably more resistant than wild type *P. fluorescens* against all the effector-delivering strains, demonstrating that it is resistant to T6SS attack rather than specifically resistant to the action of one effector (Fig. 7e,f). When the Δ*moaC* mutant was co-cultured with wild type and T6SS-inactive Db10, the slight fitness disadvantage observed in the no-attacker control above was not replicated. However recovery of the Δ*moaC* mutant was significantly higher than that of wild type *P. fluorescens* when competed with wild type but not T6SS-inactive Db10 (Fig. 7e). These data are consistent with a small but significant benefit of *moaC* deletion when exposed to T6SS attack.

### Ssp4 is very common in *Serratia* and represents a new family of widely-occurring effectors

We previously reported that homologues of Ssp6 can be found across the *Enterobacterales* but not outside this order^14^. Given the differences between Ssp4 and Ssp6, including target species specificity, we investigated the distribution of Ssp4-like proteins. Interrogation of the previously-reported pan-genome of the genus *Serratia*^33^ revealed that Ssp4 is more widely distributed across the genus and occurs much more frequently than Ssp6 (Fig. 8a), which may be a consequence of its greater efficacy (Fig. 2). Given this level of conservation in *Serratia*, and the polyphyletic pattern of occurrence across the genus, we were interested to see how widely Ssp4 is found in other bacterial species. Using HMMER homology searching, we identified Ssp4 homologues across the *Gammaproteobacteria* (Fig. 8b). Ssp4 homologues in a number of genera within the order *Enterobacterales* formed a group closely related to Ssp4 in *S. marcescens*, whilst two other groups of more distantly-related Ssp4-like proteins were observed, one containing a large number of Ssp4 homologues across the genus *Pseudomonas*, and the other containing Ssp4 homologues in several orders of marine bacteria (Fig. 8c). Examining the genomic context of these Ssp4 homologues showed that Ssp4 genes are encoded next to a small downstream gene that is likely to encode a cognate immunity protein (Fig. 8d). Additionally, for all three groups of Ssp4-like proteins, whilst the corresponding genes are often found in loci distant from any T6SS genes, in some cases they are found next to genes encoding T6SS components which recruit effectors (Hcp and VgrG) or within large T6SS gene clusters (Fig. 8d), strongly indicating that these groups of Ssp4-like proteins are also T6SS-dependent effectors. Overall, this analysis indicates that Ssp4 is the founding member of a new family of ion-selective pore-forming effectors, which contains at least three groups spanning a number of bacterial orders.

**Figure 8.**
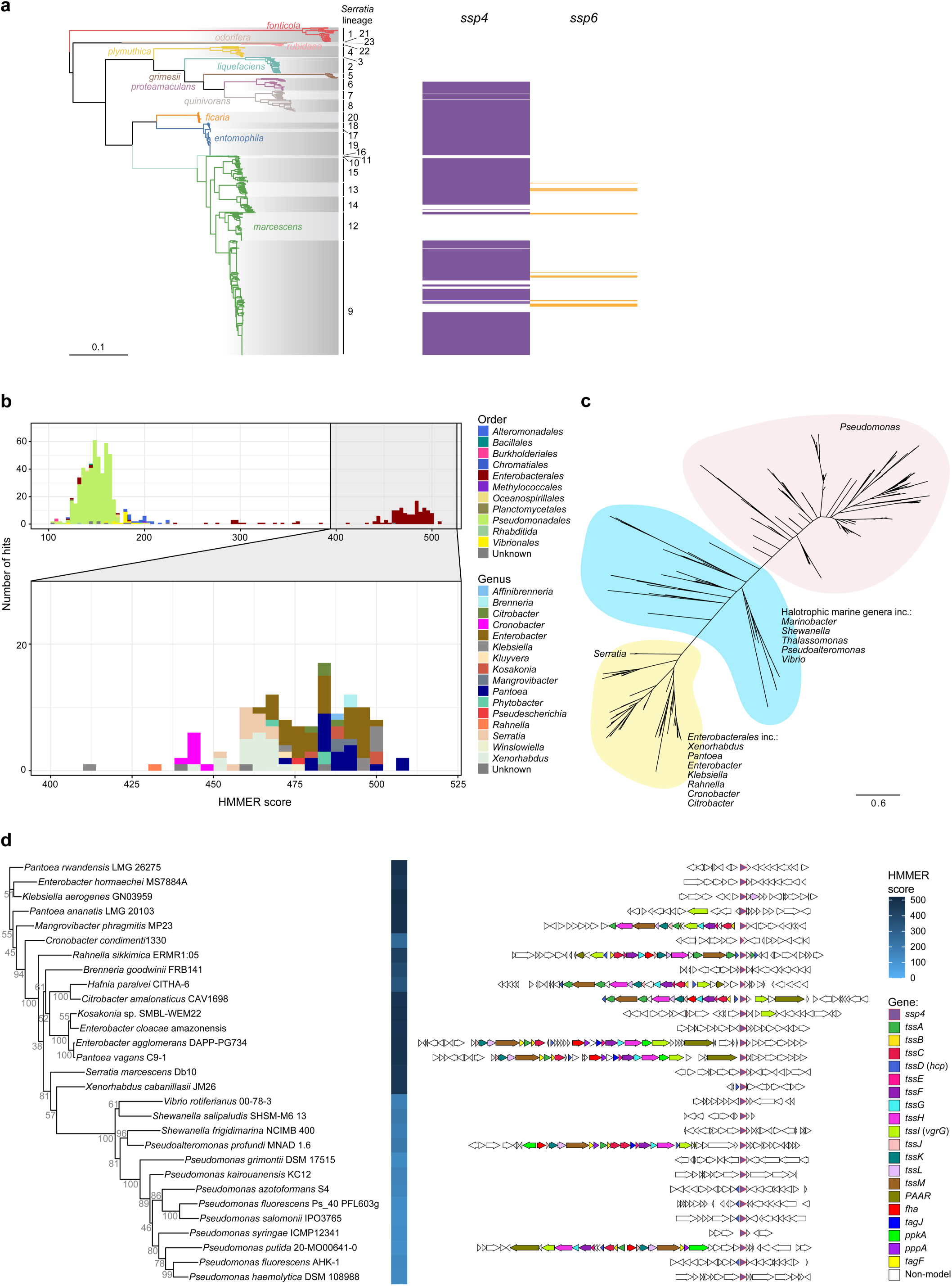
Ssp4-like proteins occur very frequently in *Serratia* and widely across other genera. (a) Presence/absence of the *ssp4* gene (purple) and the *ssp6* gene (orange) shown alongside a maximum-likelihood phylogenetic tree of 664 *Serratia* genomes determined previously from a core gene alignment (Williams et al., 2022). (b) Homologues of Ssp4 identified using HMMER homology searching, coloured by Order of the encoding bacteria (top panel) or by Genus (bottom/expanded panel). (c) Unrooted maximum-likelihood phylogeny of all identified Ssp4 homologues showing the three groups formed by Ssp4-like proteins in *Pseudomonas* (pink), marine genera (cyan), and Enterobacterales (yellow). (d) Maximum likelihood tree of selected Ssp4 protein sequences from panel c, with hmmsearch score (left) and the genetic context of the corresponding encoding gene (right). Bootstrap values are indicated on the tree and the scale indicates number of substitutions per site. Conserved T6SS genes and *ssp4* are coloured as per the key, with other genes white.

## Discussion

In this study, we have shown that Ssp4, an effector delivered by the T6SS of *S. marcescens* Db10, is a potent antibacterial toxin that forms cation-selective pores in the inner membrane of intoxicated cells, leading to a catastrophic disruption of their membrane potential. Homologues of Ssp4 are found widely in Gram-negative bacteria and represent a new family of T6SS effector proteins with at least three sub-groups.

The discovery that Ssp4 is a cation-selective pore forming effector was somewhat unexpected, given that Db10 possesses another T6SS effector, Ssp6, which, although unrelated to Ssp4, is also a cation-selective pore forming toxin and causes similar membrane depolarisation and cessation of growth in intoxicated cells. However, the two effectors are not redundant, and differ in their potency, ion selectivity, target species specificity, and the distribution of their homologues across bacterial genera. T6SS effectors are often considered to be ‘broad spectrum’ based on the fact that they target conserved, essential cellular structures and molecules in bacterial cells. However, here we observed that whilst Ssp4 was able to effectively intoxicate *S. marcescens, E. coli* and *P. fluorescens*, Ssp6 was not able to intoxicate *P. fluorescens*, implying it may only work against species relatively closely related to *Serratia*. Interestingly, whilst homologues of Ssp6 are restricted to the Enterobacterales, Ssp4-like proteins are found much more widely, including in *Pseudomonas* and *Vibrio* species. Ssp4 is also much more commonly found across *Serratia* than Ssp6. Taken together, we suggest that Ssp4-type effectors are commonly used for competition between more distantly related species, whilst Ssp6 is used only for intra-order competition within the Enterobacterales. This broader specificity, coupled with the higher potency of Ssp4, may explain why Ssp4 is found so frequently in *S. marcescens*. However, this frequency also means it is less useful for competition within species, hence certain isolates gain a competitive intra-species advantage by horizontal acquisition of the rarer Ssp6.

Structural prediction reveals that Ssp4 is an α-pore forming toxin (α-PFT), a class which includes several colicins^34^ and likely other T6SS pore-forming effectors, including Ssp6, Tse5 and Tse4 from *P. aeruginosa*, and VasX from *V. cholerae*^13, 14, 35, 36^. Upon delivery to their target cellular compartment, α-PFT monomers typically undergo a conformational change resulting in exposure of hydrophobic or amphipathic helices, followed by oligomerisation and concomitant membrane insertion to form the mature pore^34^. Such toxins are thus well suited to delivery by the T6SS, where two forms of the protein are likely to exist: a pre-secretion form which is soluble and compact to allow loading into the Hcp tube, and an active form with exposed hydrophobic helices primed for membrane insertion. The basis for the resistance to Ssp6 intoxication observed in *P. fluorescens* remains to be elucidated. While it cannot cause detectable depolarisation, some Ssp6 can be found associated with the membrane fraction. Differences in lipid composition or membrane structure might mean that pore formation cannot proceed beyond an initial interaction with the membrane and no mature pore is formed.

To date, no structures have been reported for membrane pores formed by T6SS-delivered effectors. Here, we present the first high confidence structural prediction for such a pore, supported by molecular dynamics simulations and comparison with experimental electrophysiology data. Our modelling suggests that Ssp4 forms tetrameric pores, with water-filled, ion-conducting channels formed by a total of 16 transmembrane alpha-helices. The formation of a tetrameric pore requires multiple Ssp4 proteins to be delivered into a recipient cell, most likely simultaneously. Ssp4 is an Hcp-dependent cargo effector^16^, meaning that it interacts with one or two hexameric rings of Hcp inside the T6SS-delivered puncturing structure. Given that the Hcp tube may comprise over 200 rings of Hcp^37^, it is likely that many individual Ssp4 proteins are delivered per firing event, allowing several tetrameric pores to form in the recipient cell, even from just one shot. In contrast, if Ssp4 were delivered by association with VgrG or PAAR, one shot would not deliver enough Ssp4 to form even a single pore. This example illustrates the potential importance of delivery mode to effector function.

We have demonstrated experimentally that, in common with Ssp6, Tse5, and most likely Tse4^13, 14, 35^, Ssp4 forms ion-conducting pores in target cell membranes. This leads to unregulated movement of ions across the inner membrane and disrupted membrane potential, in turn interfering with the proton motive force, ATP synthesis, membrane transport and a variety of other cellular processes^38^. These effectors cause membrane depolarisation without affecting the overall integrity of the inner membrane. In contrast, several T6SS effectors cause permeabilisation and loss of membrane integrity, suggesting they may form larger, non-specific pores, namely VasX^36^ and Tme1/2 from *V. parahaemolyticus*^39^. Whilst Ssp4 does not cause lysis or loss of membrane integrity, it can cause terminal inhibition of growth and loss of viability, as shown by the number of surviving viable target cells being much lower than the initial inoculum in co-culture assays (Fig. 2a,b).

Of the pore-forming T6SS effectors described to date, ion selectivity has only been determined *in vitro* for Ssp4, Ssp6 and Tse5, whilst cell-based assays suggested that Tse4 forms pores specific for monovalent cations^13, 14, 35^. It is interesting to note that all these effectors form pores with a strong preference for cations, although Ssp6 appears to be exclusive for cations whilst Ssp4, like Tse5, is also able to transport anions. The ability of Ssp4 to conduct anions as well as cations may contribute to its greater toxicity *in vivo*, although the ionic gradient(s) dissipated by all these toxins will vary with the extracellular environment of the intoxicated cell. Intriguingly, our data also revealed that the activity of the Ssp4 pore may be affected by the presence of Ca^2+^ at the cytoplasmic face of the inner membrane, since the properties of the pore changed when Ca^2+^ ions were present in the *trans* chamber. We predict that the *trans* chamber is equivalent to the cytoplasm in our system, since Ssp4 acts, and therefore must enter the inner membrane, from the periplasm *in vivo*, which corresponds to its addition to the lipid bilayer on the *cis* side *in vitro*. Ca^2+^ has been shown to modulate the transport properties of Tse5 and the SARS-CoV-2 E protein, suggesting that regulation by Ca^2+^ may be a common property of microbial pore-forming toxins^40, 41^.

Another interesting difference between the two pore-forming effectors deployed by *S. marcescens* Db10 is that intoxication by Ssp4 but not Ssp6 leads to the accumulation of intracellular ROS in *E. coli*. Accumulation of ROS causes damage to DNA, protein and lipids and can contribute to the lethality of antibiotics under certain conditions^24, 42^. It is possible that ROS production also contributes to the higher level of toxicity observed for Ssp4 compared with Ssp6. The reason for the difference between the effectors is not clear but might perhaps reflect the ability of Ssp4 to interfere with a greater number of ion gradients. Bacteria possess multiple protective pathways that counter ROS damage. Disruption of one of these, SoxRS, resulted in increased susceptibility of *E. coli* to T6SS attacks, suggesting that ROS generation can contribute to T6SS-mediated antibacterial activity^25^. Our observation that intoxication with Ssp4, but not Ssp5 or Ssp6, causes increased intracellular ROS suggests that ROS production may be a specific consequence of certain effectors rather than a general response to the stress induced by T6SS attack. Activation of CpxA in response to cell envelope stress has been implicated in aminoglycoside-induced ROS production and its loss can increase resistance towards such compounds^28^. Deletion of *cpxA* caused only a modest reduction in ROS accumulation following expression of sp-Ssp4 suggesting that ROS production in response to Ssp4 is largely independent of the Cpx pathway. Loss of CpxA had no impact on *E. coli* susceptibility to Ssp4 in co-culture, indicating that the Cpx stress response pathway does not significantly potentiate or protect against T6SS-delivered Ssp4, consistent with observations for TseH of *V. cholerae*^43^.

The strong Ssp4-dependent antibacterial activity observed in co-culture experiments provided an opportunity to investigate whether an unbiased approach, Tn-seq, could identify mutants resistant to a pore-forming effector. One gene whose inactivation caused a modest level of resistance to Ssp4, *moaC*, was identified, although there is a trade-off with the fact that loss of MoaC, a protein required for molybdenum cofactor (Moco) biosynthesis, has a deleterious impact on overall fitness. It is currently unclear why interruption of Moco synthesis or loss of the ability to assemble Moco-containing enzymes, which are redox-active enzymes involved in electron-transfer reactions during nitrogen, carbon, and sulphur metabolism^30^, would provide an advantage under conditions of Ssp4 intoxication. One hypothesis is that eliminating activity of these enzymes in the context of elevated oxidative stress could be beneficial. MoaC acts before the step in Moco synthesis that requires activity of two proteins outside the Moco-specific pathway (IscS and TusA), hence loss of competition from MoaC could free these enzymes to aid in repairing oxidative damage to Fe-S clusters in other proteins. However, any advantage of *moaC* inactivation will only be seen in conditions when Moco-dependent electron transfer reactions are not essential, such as during growth on LB.

More strikingly, the Tn-seq approach revealed that loss of the C-terminal domain of the anti-sigma factor MucA provides *P. fluorescens* with substantially increased protection against overall T6SS attack. Given that Ssp6 intoxication does not cause a detectable phenotype in *P. fluorescens*, the selection of *mucA* mutants in the Tn-seq comparison of Δ9 + *ssp6* vs Δ9 was unexpected. It is possible that interaction of Ssp6 with the membrane in *P. fluorescens* leads to rare or immature pores conferring a small fitness defect detectable by this assay. Alternatively, reintroduction of Ssp6 into the Δ9 mutant might increase T6SS firing rate via a checkpoint for effector loading^44^, leading to increased T6SS damage to the target cells, either from penetration or delivery of an as-yet-unidentified effector.

MucA has been extensively characterised in *P. aeruginosa* due to its role in regulating production of the extracellular polysaccharide (EPS) alginate and the fact that mutations causing C-terminal truncations of MucA lead to a mucoid (alginate overproducing) phenotype in isolates from cystic fibrosis patients^32, 45^. Production of alginate in *P. fluorescens* is also controlled by MucA and truncations of MucA lead to EPS production, in concert with other changes in gene expression consistent with its role as an anti-sigma factor^46^. Previous studies have shown that production or overproduction of EPS, either as a membrane-bound capsule or free EPS, by *V. cholerae*, *E. coli* and *Klebsiella* can provide protection against T6SS attack^43, 47, 48, 49^. This protection operates by providing a physical barrier and, in the case of free EPS, by facilitating spatial segregation of producers and bystanders away from T6SS-wielding attacker cells^48^. Thus we propose that the increased survival of the *P. fluorescens mucA*_Δ76-195_ mutant is due to increased alginate EPS production, providing both a physical barrier between attacker and target cells and promoting formation of *P. fluorescens* microcolonies which spatially segregate the cells within them from *S. marcescens* attackers. In support of this model, a *P. aeruginosa mucA* mutant was shown to exhibit enhanced microcolony formation and form more highly structured biofilms, dependent on overproduction of alginate^50^.

In conclusion, Ssp4 represents the founder member of a widely-distributed family of T6SS-dependent effectors which form ion-selective transmembrane pores. Ssp4 is one of two unrelated cation-selective pore-forming effectors delivered by the T6SS of *S. marcescens* Db10 which differ in their ion selectivity, target specificity and impact on intoxicated cells. Moreover, these two effectors are likely to be used differently, one for ‘intra-order’ competition within closely related species, the other primarily between more distantly related competitors. Co-occurrence of two such pore-forming effectors may be common (e.g Tse5 and Tse4 in *P. aeruginosa*) and these effectors likely play an important synergistic role in overall T6SS-mediated antibacterial activity, as loss of membrane potential will impair the ability of intoxicated cells to repair damage or reverse nucleotide depletion caused by other types of effectors. Conversely, production of EPS is emerging as a common means of protection against T6SS assault, with alginate production by *P. fluorescens* providing another example of this phenomenon. Overall, this study provides further evidence for the diversity of effectors and complexity of competitive inter-bacterial interactions mediated by the T6SS.

## Methods

### Bacterial strains and plasmids

Bacterial strains and plasmids used in this study are detailed in Supplementary Table 1. Strains of *S. marcescens* and *P. fluorescens* were routinely grown at 30 °C, and *E. coli* at 37 °C, in liquid LB (10 g/L tryptone, 5 g/L yeast extract, 5 g/L NaCl) or on LB plates (10 g/L tryptone, 5 g/L yeast extract, 10 g/L NaCl, 1.8 % select agar). Where required, growth media was supplemented with 100 µg/ml ampicillin (Amp), 100 µg/ml kanamycin (Kan), 50 µg/ml gentamicin (Gen), or 100 µg/ml streptomycin (Str). Defined chromosomal mutations, including in-frame deletions and restoration of wild type alleles, in *S. marcescens* Db10 and *P. fluorescens 55* were generated by allelic exchange using the plasmids pKNG101 or pMQ30, respectively^51, 52^. *S. marcescens* Db10 differs by only a single nucleotide from strain Db11 and thus the complete genome sequence of Db11 is used interchangeably for Db10^53^. Exogenous genes were integrated into the neutral *attB* site of *P. fluorescens* using the plasmids pJM220 (for rhamnose-inducible expression constructs) or pUC18T-miniTn*7*T-Gm^R^ P_rpsG_-mScarlet (for gentamycin resistant target strains)^51^. Selected mutants of *E. coli* BW25113 were retrieved from the Keio collection^29^, verified using PCR and sequencing, and if required, the kanamycin resistance cassette was excised by transient expression of FLP recombinase from the plasmid pCP20^29^. Plasmids for arabinose-inducible gene expression were derived from pBAD18-Kn^54^ and for constitutive gene expression from pSUPROM^55^. To artificially direct export of heterologously-expressed effectors to the periplasm, an OmpA signal peptide (sp; sequence MKKTAIAIAVALAGFATVAQAAPK) was incorporated at the N-terminus of the protein. Oligonucleotide primers and details of plasmid construction are in Supplementary Table 2. All bacterial strains and plasmids generated in this study are available from the corresponding author on reasonable request.

### Bacterial co-culture (competition) assays

Co-culture assays for T6SS-dependent antibacterial activity were based on the method originally described by Murdoch *et al.*^52^. Attacker and target cells were normalised to an OD_600_ of 0.5 in LB, combined at an initial ratio of 1:1 and 25 µl of the mixture spotted onto pre-warmed LB agar plates. The co-cultures were incubated at 30°C (*P. fluorescens* and *S. marcescens* targets) or 37°C (*E. coli* targets) for 4 h and then the recovery of viable target cells was enumerated by resuspending the total population in liquid media, performing serial 10-fold dilutions, and plating on media containing the appropriate antibiotic to select the target cells.

### SDS-PAGE and immunoblotting

Protein samples were combined with 4x SDS-PAGE loading buffer (200 mM Tris-HCl pH 6.8, 6.4% SDS, 6.4 mM EDTA, 32% glycerol, 0.07% bromophenol blue, 5% β-mercaptoethanol) and boiled for 5 min. 5 µl aliquots were separated on 4–20 % Mini-PROTEAN TGX Precast Protein Gels (Bio-Rad) and stained with Instant Blue (Expedeon). For immunoblotting, gels were transferred onto nitrocellulose membranes using the iBlot 2 Gel Transfer Device (Thermo), blocked with 2.5 % milk powder (Marvel) in PBS + 0.1 % Tween-20 and probed with affinity purified anti-Ssp4 (1:200) or anti-Ssp6 antibodies (1:2000). Bound antibodies were detected using a 1:10,000 dilution of HRP-conjugated goat anti-rabbit secondary antibody (BioRad #170-6515) and an enhanced chemiluminescence detection kit (Millipore). EF-Tu was detected using a 1:10,000 dilution of mouse anti-EF-Tu antibody (Hycult #HM6010), the 3xFLAG epitope tag was detected using a 1:10,00 dilution of mouse anti-FLAG (Sigma, #F3165), with an HRP-conjugated goat anti-mouse secondary antibody (BioRad #170-6516). Signal detection was performed using the Azure 600 imaging system (Cambridge Bioscience).

### Antiserum generation

For custom antiserum generation, *E. coli* BL21(DE3) was transformed with pET15b-based plasmids directing the expression of TEV-cleavable N-terminal His_10_-tagged Ssp4 or Ssp6. Cultures in 500 ml LB were induced with 0.5 mM IPTG overnight at 16°C and recombinant Ssp4 and Ssp6 were purified by Ni-NTA chromatography under denaturing conditions according to the methodology originally described in^56^. Cells were resuspended in 1/25^th^ volume denaturing lysis buffer (20 mM sodium phosphate pH 7.8, 500 mM NaCl, 20 mM imidazole, 1% SDS), lysed by sonication and incubated with 1 ml of settled Ni-NTA agarose for 1 h at room temperature. The bound protein was washed extensively with denaturing wash buffer (20 mM sodium phosphate pH 7.8, 500 mM NaCl, 50 mM imidazole, 0.1% sarkosyl) and eluted in 8x 1 ml fractions in denaturing elution buffer (20 mM sodium phosphate pH 7.8, 500 mM NaCl, 250 mM imidazole, 0.1% sarkosyl). Eluates were combined with 4x SDS-PAGE loading buffer and 10 µl aliquots were separated using 4–20 % Mini-PROTEAN TGX Precast Protein Gels and stained with InstantBlue. Ssp4 or Ssp6 bands were excised for protein extraction and rabbit antiserum generation (Davids Biotechnologie).

### Antibody affinity purification

Recombinant untagged Ssp4 and His-GST tagged Ssp6 (prepared as below) were coupled to cyanogen bromide activated agarose (Sigma Aldrich) based on the method described in^57^. Proteins were dialysed into an appropriate volume of coupling buffer (0.1 M NaHCO_3_ pH 8.3, 0.5 M NaCl) and incubated with 1 ml of swollen cyanogen bromide activated agarose at 4°C overnight. Remaining active sites were blocked with 0.2 M glycine (1 h, room temperature) and then the resin was washed extensively with coupling buffer followed by low pH buffer (0.1 M C_2_H_3_NaO_2_ pH 4.0, 0.5 M NaCl). The high/low pH buffer wash cycle was repeated for a total of five times and then the affinity purification resin was equilibrated into TBS.

For the antibody purification, 1 ml aliquots of rabbit antiserum were diluted 1:5 in TBS and incubated with the affinity purification resin for 2 h at room temperature. Antibodies were eluted in 8 ml of 1 M acetic acid which was immediately neutralised using an equal volume of 2.5 M Tris-HCl pH 8.8. Purified antibodies were exchanged into TBS using a HiPrep 26/10 Desalting column (Cytiva) and concentrated to approximately 1 ml using a 50,000 MWCO centrifugal filter (Amicon). The purified antibodies were supplemented with 0.02% BSA and 0.02% sodium azide and stored at 4°C. To further deplete non-specific antibodies from the purified anti-Ssp6 preparation, the blocking buffer was pre-incubated with solubilised wild type *P. fluorescens* membranes (prepared as below) at a final concentration of 1:10,000 and for 2 h prior to addition to the nitrocellulose membrane.

### PEG Labelling of periplasmic/cytoplasmic cysteines

Free cysteines were labelled using methoxypolyethylene glycol maleimide (mPEG-MAL, Sigma Aldrich). Cultures of *S. marcescens* were grown in LB at 30°C for 5 h. Cells were collected by centrifugation at 4,000 x*g* for 10 min, washed once with HEPES/MgCl_2_ buffer (50 mM HEPES pH 6.8, 5 mM MgCl_2_) and resuspended in HEPES/MgCl_2_ buffer to a final cell density of 0.3 OD units/ml. 80 µl of cell suspension was incubated with 5 mM mPEG-MAL and 10 mM EDTA in a final rection volume of 100 µl HEPES/MgCl_2_ buffer for 1 h at room temperature. Duplicate labelling reactions were performed in the presence of 1% SDS to disrupt the cell envelope and allow mPEG-MAL labelling of transmembrane and cytoplasmic cysteine residues. The labelling reaction was stopped by addition of DTT (final concentration 100 mM) and the labelled cells were collected by centrifugation at 21,000 x*g* for 5 min and resuspended in 100 µl of 1x SDS-PAGE loading buffer.

### Fractionation of *P. fluorescens* cells

Cultures of *P. fluorescens* were grown in 15 ml LB at 30°C for 3 h and induced with 0.05% L-rhamnose for 2 h at 30°C. Cells were collected by centrifugation at 4000 x*g*, resuspended in 1.5 ml of sucrose buffer (10 mM HEPES pH 7.4, 10% sucrose, 10 mM EDTA, 150 µg/ml lysozyme, 50 U/ml Benzonase nuclease) with protease inhibitors (cOmplete EDTA-free Protease Inhibitor Cocktail, Roche) and incubated on ice overnight. Whole cell lysates were prepared by sonication using a 2 mm probe (Amplitude 20%; 6x cycles of 15s on/off) and removal of cell debris by two rounds of centrifugation at 20, 000 x*g*. Soluble and membrane associated fractions were separated by ultracentrifugation at 50,000 rpm (TLA-120.2 Fixed-Angle Rotor: R_av_ ∼89,000 x*g*) for 30 min at 4°C. Membrane pellets were washed twice with 10 mM HEPES pH 7.4 prior to solubilisation in 200 µl 2x SDS-PAGE loading buffer.

### Microscopy

Strains of *S. marcescens* were grown in 25 ml minimal glucose media (40 mM K_2_HPO_4_, 15 mM KH_2_PO_4_, 0.1% (NH_4_)_2_SO_4_, 0.4 mM MgSO_4_, 0.2% w/v glucose) supplemented with 50 µM IPTG at 30°C for 4.5 h and adjusted to an OD_600_ of 0.5. Attacker and target cells were combined in an initial 3:1 ration and 1 µl aliquots of the mixture were spotted onto a microscope slide layered with a pad of minimal glucose medium + 50 µM IPTG solidified by the addition of 1.5% UltraPure agarose (Invitrogen) and sealed with 1.5 thickness coverslips (VWR) attached to the microscope slide with a GeneFrame (ThermoFisher Scientific). The slides were allowed to equilibrate within the microscope chamber, pre-heated to 30°C, for approx. 45 minutes during which time 10-15 frames containing mixed attacker-target microcolonies were selected. Imaging was performed using a DeltaVision Core widefield microscope mounted on an Olympus IX71 inverted stand with an Olympus 100X 1.35 NA objective and Cascade2 EMCCD camera. Image stacks were acquired every 12 min with Z spacing of 0.3 µm. GFP (target cells) was imaged using Ex/Em 480 nm/525 nm and exposure time 100 ms, and mCherry (attacker cells) was imaged using Ex/Em 575 nm/628 nm and exposure time 150 ms. Post-acquisition, images were stored and processed using OMERO software (http://openmicroscopy.org)^58^. As fluorescence intensity values are not relevant to the analysis of cell numbers and distribution, images are presented following manual adjustment across the timecourse for clarity.

### Measurement of membrane potential and permeability and ROS levels using flow cytometry

Membrane potential and permeability assays were carried out using the methodology described by Mariano *et al.*^14^ with minor modification. Co-cultures between strains of *S. marcescens* were performed as described above with an initial attacker:target ratio of 1:2. Following the 4 h co-incubation, cells were collected into 1 ml of PBS, adjusted to 1x10^6^ cell/ml and stained with 10 μM DiBAC_4_(3) and 1 μM propidium iodide for 30 min on ice in the dark. For analysis of the impact of sp-Ssp4 and sp-Ssp6 expressed heterologously in *P. fluorescens*, overnight cultures were diluted 1 in 25 in LB and incubated at 30°C for 3 h, gene expression was induced by the addition of 0.05% L-rhamnose and the cultures incubated for a further 2 h prior to staining with DiBAC_4_(3) and PI as above. Controls were prepared using exponential phase cultures of *P. fluorescens* that had been treated with 6.3 µM melittin (Sigma Aldrich) in PBS for 2 h at 30°C.

For detection of ROS, overnight cultures of *E. coli* MG1655 or BW25113 carrying pBAD18-Kn derived plasmids were diluted 1 in 100 into LB media containing 10 µM OxyBURST^TM^ Green (2’,7’-dichlorodihydrofluorescein diacetate, succinimidyl ester; Invitrogen #D2935) and incubated at 37°C for 2 h. Gene expression was induced by addition of 0.2 % L-arabinose and incubation continued for a further 3 h. At each time point at or post-induction, 10 µl of induced culture was fixed with 1% formaldehyde in PBS for 30 min at 4°C.

Samples were analysed using a LSRFortessa Cell Analyzer (BD) equipped with 488 nm and 561 nm lasers. Data analysis was performed using FlowJo v10.4.2 (Treestar). Gating strategies are provided in the Supplementary Information (Suppl. Fig. 3). 20,000 and 10,000 P1 events were collected in DiBAC_4_(3)/PI and OxyBURST experiments, respectively

### Recombinant Ssp4 production

For production of His_6_-GST-Ssp4 and His_6_-GST-Ssp4, *E. coli* SHuffle T7 was transformed with plasmids derived from a modified version of the commercial pGEX-6-P1 vector (Suppl. Table 1). Cultures were grown to OD_600_ ∼ 0.6-0.8 in 6 L LB at 30°C, then gene expression was induced by the addition of 0.5 mM IPTG prior to overnight growth at 16°C. Cells were collected by centrifugation and resuspended in 150 ml of lysis buffer (50 mM Tris-HCl pH 7.5, 0.5 M NaCl, 20 mM imidazole, 2 mM TCEP) supplemented with complete EDTA-free protease inhibitor cocktail (Millipore) and 50 U/ml Benzonase nuclease (Millipore). Cells were broken by pressure cell lysis at 25,000 psi and the lysate clarified by centrifugation at 48,000 x*g* and passage through a 0.45 µm filter. Recombinant proteins were isolated by immobilised metal affinity chromatography using a three-step isocratic elution (60 mM, 120 mM and 200 mM imidazole) and a 5 ml HisTrap HP column (Cytiva). Protein-containing fractions were pooled and incubated with 1 ml of glutathione sepharose 4B resin (Cytiva) for 1 h at room temperature. The resin was washed with 20 ml of GST buffer (20 mM Tris-HCl pH 7.5, 0.5 M NaCl, 1 mM TCEP, 1 mM EDTA) and His-GST tagged proteins were eluted in 5 ml GST buffer containing 50 mM reduced glutathione.

To obtain untagged Ssp4, samples were exchanged into buffer containing 20 mM Tris-HCl pH 7.5, 0.5 M NaCl, 1 mM TCEP using a HiPrep 26/10 Desalting column and incubated with 10 µl of PreScission protease (Cytiva) overnight at 4°C. Following removal of the cleaved His_6_-GST using a 1:1 mixture of glutathione and Ni-NTA sepharose (100 µl each, 1 h incubation at room temperature), recombinant Ssp4 was concentrated using a 30,000 MWCO centrifugal filter and further purified by size exclusion chromatography (SEC) using a HiLoad S200 16/600 column (Cytiva) in buffer containing 20 mM Tris-HCl pH 7.5, 0.5 M NaCl, 1 mM TCEP. The folded state and stability of the protein was assessed by SEC using a calibrated HiLoad S200 16/600 column and thermal calorimetry using the Tycho NT.6 system (NanoTemper).

### Electrophysiology measurements and analysis

Planar lipid bilayers were prepared by resuspending bovine phosphatidylethanolamine lipids (Avanti Polar Lipids) in decane at a final concentration of 30 mg/mL and forming lipid bilayers across a 150 μm diameter aperture in a partition that separates two 1 mL compartments, the *cis* and the *trans* chambers^59^. KCl or CaCl_2_ solutions at the concentrations indicated were buffered with 10 mM HEPES pH 7.2 and added to the appropriate chamber.

Ssp4 (0.5-1 μg) was added to the *cis* chamber. The *cis* side was continuously stirred to facilitate incorporation of Ssp4 into the bilayer and incorporation was assessed by visualisation of channel activity measured by a change in current from 0 pA. Following Ssp4 incorporation, the *trans* chamber was held at 0 mV while the *cis* chamber was clamped at different holding potentials relative to ground. The transmembrane current was measured under voltage-clamp conditions using a BC-525C amplifier (Warner Instruments, Harvard Instruments). Channel recordings were low-pass filtered at 10 kHz with a four-pole Bessel filter, digitized at 100 kHz using a National Instruments acquisition interface (NIDAQ-MX, National Instruments, Austin, TX) and recorded on a computer hard drive using WinEDR 4.00 acquisition software (John Dempster, University of Strathclyde, Glasgow, UK). Current fluctuations were measured at room temperature over 30-60 s. Recordings were filtered using a low pass digital filter at 800 Hz (-3dB) implemented in WinEDR 3.05. Measurements of current amplitudes were carried out in WinEDR 3.05. Predicted reversal potentials were calculated using the Nernst equation. The reversal potential (*E_rev_*) was taken at the voltage where zero current was measured. Junction potentials were calculated using Clampex v10.2 (Molecular Devices) and subtracted from the reversal potential obtained for each experiment.

### Generation of saturated *P. fluorescens* 55 transposon mutant library

*E. coli* SM10λpir carrying pIT2 and *P. fluorescens* 55 were each streaked onto three LB + Amp (*E. coli*) or LB (*P. fluorescens*) plates from frozen stocks and grown overnight. Cells were recovered and resuspended to an OD_600_ of 50 (*P. fluorescens*) or 100 (*E. coli*) in LB. 100 µl of each suspension were combined and 50 µl aliquots were spotted onto LB agar followed by incubation at 30°C for ∼6 h. Cells were resuspended in LB, diluted and plated onto ∼60 LB agar plates containing 60 µg/ml tetracycline (to select for *P. fluorescens* cells containing transposon insertions) and 10 µg/ml chloramphenicol (to counterselect *E. coli*) and incubated at 30°C for 24 h. Resulting colonies (∼60,000 – 100,000) were resuspended in 6 ml LB to which 3 ml of 50% glycerol was added for storage at -80°C in 100 µl aliquots.

### Co-culture and library preparation for transposon insertion site sequencing

Similar to the co-culture assays above, cells of each strain of *S*. *marcescens* and an aliquot of the *P. fluorescens* library were adjusted to an OD_600_ of 0.5 in LB. The suspensions were combined 1:2 (attacker:target) in appropriate combinations, and 2x 25 µl aliquots of the mixture were spotted onto pre-warmed LB agar plates and incubated at 30°C for 4 h. The resulting cells were resuspended in 1 ml of PBS and genomic DNA was extracted using the DNeasy Blood & Tissue Kit (Qiagen). Genomic DNA (∼10 µg) was sheared into 250 bp fragments by sonication using the QSonica Q800R sonicator (Amplitude 25%: 20x cycles of 15s on/off) and end repaired using the NEBNext End Repair Module (New England Biolabs). A polyC tail was added to 1 µg of end repaired DNA using Terminal Deoxynucleotidyl Transferase (NEB) with a mixture of 95% dCTP and 5% ddCTP as a substrate. Residual C-tailing reagents were removed using NucleoSeq gel filtration columns (Macherey-Nagel) and DNA fragments containing transposon insertion sites were amplified by PCR using Q5 Hot Start High-Fidelity DNA Polymerase (New England Biolabs) and oligonucleotide primers as detailed in Supplementary Table 2. Sample clean-up and size selection was performed using magnetic AMPure XP beads (Beckman). An initial incubation with 0.8 volumes of bead solution was used to remove long DNA fragments (>250 bp) and the fragments of interest were adsorbed from the resulting supernatant using an additional 0.4 volumes of bead solution. The beads were washed twice with 80% ethanol prior to elution of the DNA in 30 µl ultrapure H_2_O. Index primers (NEBNext Multiplex Oligos for Illumina, New England Biolabs) were incorporated by PCR and indexed fragments of interest were adsorbed using 1.2 volumes of AMPure XP beads. The beads were washed twice with 80% ethanol prior to elution of the DNA in 30 µl ultrapure H_2_O. The quality and concentration of DNA was assessed using High Sensitivity DNA ScreenTape (Agilent) and an Agilent 2200 TapeStation.

### Generation and analysis of Tn-seq sequencing data

Prepared DNA libraries were pooled for sequencing on the Illumina NextSeq2000 instrument with a P1 reagent kit. Approximately 2.2 - 4.6 million single-end 150 bp reads were obtained per sample. Raw FASTQ files were processed for analysis using the Tn-Seq Pre-Processor (TPP) tool from the Transit package^60^, which finds and filters the transposon sequence from each read and maps the remaining genome sequence to a reference genome (here *P. fluorescens* 55, see below) using bwa (http://arxiv.org/abs/1303.3997). Approximately 1.9 - 4.0 reads per sample were successfully mapped. Plots of insertions per site were generated from .wig files output by the TPP tool and output .sam files were used in conjunction with the genome annotation to summarise counts per gene using the featureCounts algorithm of the subread software package^61^.

We aimed to identify genes with significantly altered numbers of transposon-derived reads between conditions. Because the fate of individual cells in a T6SS intoxication experiment is stochastic (depending on proximity of each prey cell to an attacker cell, for example) we found that in many cases the normalised counts per gene varied dramatically between the three biological replicates of each intoxication condition. We assume that these highly variable genes do not strongly affect fitness and therefore the dataset for subsequent analysis was limited to genes where the coefficient of variation under each condition was < 0.5. Differential expression analysis was then performed with *edgeR* version 3.38.4^62^ to compare pairwise the three intoxication conditions (Δ9 vs Δ9 + *ssp4*; Δ9 vs Δ9 + *ssp6*; Δ9 + *ssp4* vs Δ9 + *ssp6*) in order to to identify genes significantly different between condition (FDR < 0.05). The R code for this analysis is available at https://github.com/bartongroup/MG_T6SS_tn-seq.

Separately, whole genome sequencing of *P. fluorescens* 55 was provided by MicrobesNG (https://microbesng.com), using using hybrid short (Illumina) and long (Oxford Nanopore) read sequencing to generate a closed whole genome assembly with automated gene annotation.

### Structural predictions

Membrane topology predictions were generated using MEMSAT-SVM^63^ hosted on the PSIPred workbench (http://bioinf.cs.ucl.ac.uk/psipred/). Predictions of Ssp4 and Sip4 monomer structures were generated using AlphaFold2^12^ hosted on Google Colab (https://colab.research.google.com). Structural predictions for dimeric, trimeric, tetrameric, hexameric and octameric Ssp4 assemblies (putative pore structures) were generated using AlphaFold2^12^ hosted on the University of Dundee HPC cluster and propensity for membrane insertion was predicted using MemProtMD^21^ with INSANE (https://github.com/pstansfeld/MemProtMD) hosted on GoogleColab. Structural representations were generated using the PyMOL Molecular Graphics System, Version 2.0 (Schrödinger).

### Molecular Dynamics simulations

The structural model of an Ssp4_114-302_ tetramer was embedded into a 1-palmitoyl-2-oleoyl-sn-glycerol-3-phosphatidyl choline (POPC) membrane and aligned in the membrane with PPM^64^ using the CHARMM-GUI server^65^. The initial box size was 90 x 90 Å in x,y-dimension and 110 Å in z-dimension. The CHARMM36m force field was used for the protein, lipids and ions (KCl at a concentration of 0.15M), and the TIP3P water model was used to model water molecules^66, 67^. All Molecular Dynamics (MD) simulations were carried out with the GROMACS 2022 software package^68^. Energy minimization and equilibration was performed according to the protocols provided by the CHARMM-GUI server^65^. A further equilibration step comprised unbiased simulations of 10 ns length without restraints or an electric field, using a 1 fs integration timestep. The stability of the tetramer in membranes was initially studied in three-fold replicated production simulations of 300 ns length, followed by three-fold replicated production simulations under membrane voltage of 1 µs length each. The production simulations were conducted in the NPT ensemble, with the temperature controlled at 310 K using the Nosé-Hoover thermostat and the pressure semi-isotropically maintained at 1 bar using the Parrinello-Rahman barostat^69, 70^. All production simulations used a 2 fs integration timestep. To constrain bond lengths involving H atoms, the LINCS algorithm was employed; long-range electrostatic interactions were modelled using the Particle-Mesh Ewald method^71, 72^. Membrane voltages were generated using an applied external electric field^73^.

### Identification of Ssp4-like effector family

Distribution of Ssp4-like effectors across known protein space was determined by searching the UniProt database^74^ for Ssp4-like proteins, using hmmsearch from the HMMER suite v3.1b2^75^. An “Ssp4” model was constructed, using hmmbuild, from a small, manually-curated alignment of non-redundant ssp4 homologues identified using BLASTp^76^. Accessions for publicly-available genomes or contigs corresponding with Ssp4-like UniProt IDs identified using this approach were located using the ID-mapping tool in UniProt (https://www.uniprot.org/id-mapping), matching the UniProt IDs against the target database EMBL/GenBank/DDBJ. Annotated genomes corresponding to these accessions were then downloaded using the NCBI batch entrez tool (https://www.ncbi.nlm.nih.gov/sites/batchentrez). Genomic regions of ∼20 kb containing the gene encoding the Ssp4-like protein (10 kb upstream and downstream of the Ssp4-encoding gene) were then retrieved from these genomes using hamburger^14, 77^. T6SS genes were also identified during the search, using models within hamburger^14, 77^. T3SS genes were identified by hmmsearch^75^ using models from macsyfinder^78^. Several potential Ssp4 hits with a low hmmsearch sequence score matched the Type III Secretion System (T3SS) translocon protein SipB, suggesting that they were false positives. Subsequently, all hmmsearch hits with a sequence score of less than 100 were removed from further analysis. Trees of Ssp4-like protein sequences were drawn using IQTREE (v1.6.5)^79^ with 1000 ultrafast bootstraps^80^ using models chosen by modelfinder^81^. Trees were drawn from alignments created by hmmsearch^75^. Trees were visualised using the R packages ggtree (v1.15.6)^82^ and figtree (v1.4.4) (http://tree.bio.ed.ac.uk/software/figtree/), and associated genomic context depicted using ggplot2 (v3.1.1)^83^ and gggenes (v0.3.2) (https://wilkox.org/gggenes/).

## Data availability

The complete genome sequence of *P. fluorescens* 55 and the Tn-seq sequencing data will be made publicly available in the NCBI Genome and GEO databases, respectively, upon publication. The R code for the differential expression analysis is available at https://github.com/bartongroup/MG_T6SS_tn-seq. All other data supporting the findings of this study are available within the paper and its supplementary information files.

## Acknowledgements

This work was supported by Wellcome (grant numbers 104556/Z/14/Z, Senior Research Fellowship S.J.C.; and 220321/Z/20/Z, Senior Research Fellowship Renewal S.J.C.), UKRI Medical Research Council (grant numbers MR/K000111X/1, New Investigator Research Grant, S.J.C.; and MR/T041811/1, Future Leaders Fellowship, M.B.), UKRI Biotechnology and Biological Sciences Research Council (grant numbers BB/M010996/1 and BB/T00875X/1, EASTBio PhD studentships K.M. and A.T.G.), Academy of Medical Sciences (grant number SBF005/1096, Springboard Grant, M.B.; the Springboard scheme is funded by Wellcome, the Department of Business, Energy, and Industrial Strategy UK, the British Heart Foundation, and Diabetes UK) and the British Heart Foundation (grant number FS/PhD/22/2932, non-clinical PhD studentship Q.W.H.). We thank Yi-Chia Liu and Giuseppina Mariano for generation of strains; Daan van Aalten for the gift of pHis-GEX-6P-1; Laura Monlezun for establishing purification of GST-fused Ssp4; Pete Hedley for assistance with Tn-seq sequencing; and the Flow Cytometry and Cell Sorting Facility, High Performance Computing Facility and Dundee Imaging Facility at the University of Dundee for access to facilities and expert assistance. For the purpose of Open Access, the authors have applied a CC BY public copyright licence to any Author Accepted Manuscript version arising from this submission.

## Author Contributions

M.R. and S.J.C. conceived the study; M.R., Q.W.H., K.M., A.O. and M.B. performed experimental work; M.R., D.J.W., A.T. and U.Z. performed bioinformatics and computational analyses; M.R., Q.W.H., M.G., A.T., M.B., U.Z., S.J.P and S.J.C analysed data; M.R. and S.J.C. wrote the manuscript with input from D.J.W., M.B. and the other authors.

## References

1. Granato ET, Meiller-Legrand TA, Foster KR. The Evolution and Ecology of Bacterial Warfare. Curr Biol 29, R521–R537 (2019).

2. Peterson SB, Bertolli SK, Mougous JD. The Central Role of Interbacterial Antagonism in Bacterial Life. Curr Biol 30, R1203–R1214 (2020).

3. Coulthurst S. The Type VI secretion system: a versatile bacterial weapon. Microbiology, (2019).

4. Allsopp LP, Bernal P, Nolan LM, Filloux A. Causalities of war: The connection between type VI secretion system and microbiota. Cell Microbiol 22, e13153 (2020).

5. Gallegos-Monterrosa R, Coulthurst SJ. The ecological impact of a bacterial weapon: microbial interactions and the Type VI secretion system. FEMS Microbiol Rev 45, (2021).

6. Robitaille S, et al. Community composition and the environment modulate the population dynamics of type VI secretion in human gut bacteria. Nat Ecol Evol 7, 2092–2107 (2023).

7. Jurenas D, Journet L. Activity, delivery, and diversity of Type VI secretion effectors. Mol Microbiol 115, 383–394 (2021).

8. Wang J, Brodmann M, Basler M. Assembly and Subcellular Localization of Bacterial Type VI Secretion Systems. Annu Rev Microbiol 73, 621–638 (2019).

9. Hood RD, et al. A type VI secretion system of Pseudomonas aeruginosa targets a toxin to bacteria. Cell Host Microbe 7, 25–37 (2010).

10. Bullen NP, et al. An ADP-ribosyltransferase toxin kills bacterial cells by modifying structured non-coding RNAs. Mol Cell 82, 3484–3498 e3411 (2022).

11. Hernandez RE, Gallegos-Monterrosa R, Coulthurst SJ. Type VI secretion system effector proteins: Effective weapons for bacterial competitiveness. Cell Microbiol 22, e13241 (2020).

12. Jumper J, et al. Highly accurate protein structure prediction with AlphaFold. Nature 596, 583–589 (2021).

13. Gonzalez-Magana A, et al. The *P. aeruginosa* effector Tse5 forms membrane pores disrupting the membrane potential of intoxicated bacteria. Commun Biol 5, 1189 (2022).

14. Mariano G, et al. A family of Type VI secretion system effector proteins that form ion-selective pores. Nature communications 10, 5484 (2019).

15. Mahlen SD. *Serratia* infections: from military experiments to current practice. Clin Microbiol Rev 24, 755–791 (2011).

16. Cianfanelli FR, Alcoforado Diniz J, Guo M, De Cesare V, Trost M, Coulthurst SJ. VgrG and PAAR Proteins Define Distinct Versions of a Functional Type VI Secretion System. PLoS Pathog 12, e1005735 (2016).

17. Fritsch MJ, Trunk K, Diniz JA, Guo M, Trost M, Coulthurst SJ. Proteomic Identification of Novel Secreted Antibacterial Toxins of the *Serratia marcescens* Type VI Secretion System. Mol Cell Proteomics 12, 2735–2749 (2013).

18. Hagan M, et al. Rhs NADase effectors and their immunity proteins are exchangeable mediators of inter-bacterial competition in *Serratia*. Nature Communications 14, 6061 (2023).

19. Srikannathasan V, et al. Structural basis for Type VI secreted peptidoglycan DL-endopeptidase function, specificity and neutralization in *Serratia marcescens*. Acta Crystallogr D 69, 2468–2482 (2013).

20. Trunk K, et al. The type VI secretion system deploys antifungal effectors against microbial competitors. Nature Microbiology 3, 920–931 (2018).

21. Stansfeld PJ, et al. MemProtMD: Automated Insertion of Membrane Protein Structures into Explicit Lipid Membranes. Structure 23, 1350–1361 (2015).

22. Kohanski MA, Dwyer DJ, Hayete B, Lawrence CA, Collins JJ. A common mechanism of cellular death induced by bactericidal antibiotics. Cell 130, 797–810 (2007).

23. Zhao X, Hong Y, Drlica K. Moving forward with reactive oxygen species involvement in antimicrobial lethality. J Antimicrob Chemother 70, 639–642 (2015).

24. Hong Y, Zeng J, Wang X, Drlica K, Zhao X. Post-stress bacterial cell death mediated by reactive oxygen species. Proc Natl Acad Sci U S A 116, 10064–10071 (2019).

25. Dong TG, Dong S, Catalano C, Moore R, Liang X, Mekalanos JJ. Generation of reactive oxygen species by lethal attacks from competing microbes. Proc Natl Acad Sci U S A 112, 2181–2186 (2015).

26. Mahoney TF, Silhavy TJ. The Cpx stress response confers resistance to some, but not all, bactericidal antibiotics. J Bacteriol 195, 1869–1874 (2013).

27. Cudic E, Surmann K, Panasia G, Hammer E, Hunke S. The role of the two-component systems Cpx and Arc in protein alterations upon gentamicin treatment in *Escherichia coli*. BMC Microbiol 17, 197 (2017).

28. Kohanski MA, Dwyer DJ, Wierzbowski J, Cottarel G, Collins JJ. Mistranslation of membrane proteins and two-component system activation trigger antibiotic-mediated cell death. Cell 135, 679–690 (2008).

29. Baba T, et al. Construction of *Escherichia coli* K-12 in-frame, single-gene knockout mutants:the Keio collection. Mol Syst Biol 2, 2006 0008 (2006).

30. Leimkuhler S, Iobbi-Nivol C. Bacterial molybdoenzymes: old enzymes for new purposes. FEMS Microbiol Rev 40, 1–18 (2016).

31. Zhang Y, et al. Structure-guided disruption of the pseudopilus tip complex inhibits the Type II secretion in *Pseudomonas aeruginosa*. PLoS Pathog 14, e1007343 (2018).

32. Schofield MC, et al. The anti-sigma factor MucA is required for viability in *Pseudomonas aeruginosa*. Mol Microbiol 116, 550–563 (2021).

33. Williams DJ, et al. The genus *Serratia* revisited by genomics. Nature Communications 13, 5195 (2022).

34. Ulhuq FR, Mariano G. Bacterial pore-forming toxins. Microbiology (Reading) 168, (2022).

35. LaCourse KD, Peterson SB, Kulasekara HD, Radey MC, Kim J, Mougous JD. Conditional toxicity and synergy drive diversity among antibacterial effectors. Nature microbiology 3, 440–446 (2018).

36. Miyata ST, Unterweger D, Rudko SP, Pukatzki S. Dual expression profile of type VI secretion system immunity genes protects pandemic *Vibrio cholerae*. PLoS Pathog 9, e1003752 (2013).

37. Wang J, et al. Cryo-EM structure of the extended type VI secretion system sheath-tube complex. Nature Microbiology 2, 1507–1512 (2017).

38. Benarroch JM, Asally M. The Microbiologist’s Guide to Membrane Potential Dynamics*Trends Microbiol* **28**, 304–314 (2020).

39. Fridman CM, Keppel K, Gerlic M, Bosis E, Salomon D. A comparative genomics methodology reveals a widespread family of membrane-disrupting T6SS effectors. Nat Commun 11, 1085 (2020).

40. Rojas-Palomino J, González-Magaña A, Queralt-Martin M, Albesa-Jové D, Alcaraz A. Proteolipidic assembly and function of the pore-forming toxin Tse5, an effector from the *Pseudomonas aeruginosa*. Biophysical Journal 123, 376a–377a (2024).

41. Lysbeth HA, et al. The SARS-CoV-2 envelope (E) protein forms a calcium- and voltage-activated calcium channel. bioRxiv, 2022.2010.2011.511775 (2022).

42. Dwyer DJ, et al. Antibiotics induce redox-related physiological alterations as part of their lethality. Proc Natl Acad Sci U S A 111, E2100–2109 (2014).

43. Hersch SJ, et al. Envelope stress responses defend against type six secretion system attacks independently of immunity proteins. Nature microbiology 5, 706–714 (2020).

44. Liang X, Kamal F, Pei TT, Xu P, Mekalanos JJ, Dong TG. An onboard checking mechanism ensures effector delivery of the type VI secretion system in *Vibrio cholerae*. Proc Natl Acad Sci U S A 116, 23292–23298 (2019).

45. Boucher JC, Yu H, Mudd MH, Deretic V. Mucoid *Pseudomonas aeruginosa* in cystic fibrosis: characterization of muc mutations in clinical isolates and analysis of clearance in a mouse model of respiratory infection. Infection and Immunity 65, 3838–3846 (1997).

46. Borgos SE, et al. Mapping global effects of the anti-sigma factor MucA in *Pseudomonas fluorescens* SBW25 through genome-scale metabolic modeling. BMC Syst Biol 7, 19 (2013).

47. Flaugnatti N, et al. Human commensal gut Proteobacteria withstand type VI secretion attacks through immunity protein-independent mechanisms. Nature communications 12, 5751 (2021).

48. Granato ET, Smith WPJ, Foster KR. Collective protection against the type VI secretion system in bacteria. The ISME journal 17, 1052–1062 (2023).

49. Toska J, Ho BT, Mekalanos JJ. Exopolysaccharide protects *Vibrio cholerae* from exogenous attacks by the type 6 secretion system. Proc Natl Acad Sci U S A 115, 7997–8002 (2018).

50. Hentzer M, et al. Alginate overproduction affects *Pseudomonas aeruginosa* biofilm structure and function. J Bacteriol 183, 5395–5401 (2001).

51. Choi KH, Schweizer HP. mini-Tn7 insertion in bacteria with secondary, non-glmS-linked attTn7 sites: example *Proteus mirabilis* HI4320. Nat Protoc 1, 170–178 (2006).

52. Murdoch SL, Trunk K, English G, Fritsch MJ, Pourkarimi E, Coulthurst SJ. The opportunistic pathogen *Serratia marcescens* utilizes type VI secretion to target bacterial competitors. J Bacteriol 193, 6057–6069 (2011).

53. Iguchi A, et al. Genome evolution and plasticity of *Serratia marcescens*, an important multidrug-resistant nosocomial pathogen. Genome biology and evolution 6, 2096–2110 (2014).

54. Guzman LM, Belin D, Carson MJ, Beckwith J. Tight regulation, modulation, and high-level expression by vectors containing the arabinose PBAD promoter. Journal of bacteriology 177, 4121–4130 (1995).

55. Jack RL, Buchanan G, Dubini A, Hatzixanthis K, Palmer T, Sargent F. Coordinating assembly and export of complex bacterial proteins. Embo J 23, 3962–3972 (2004).

56. Schlager B, Straessle A, Hafen E. Use of anionic denaturing detergents to purify insoluble proteins after overexpression. BMC Biotechnol 12, 95 (2012).

57. Reglinski M. Purification of Prospective Vaccine Antigens from Gram-Positive Pathogens by Immunoprecipitation. Methods Mol Biol 2414, 37–45 (2022).

58. Allan C, et al. OMERO: flexible, model-driven data management for experimental biology.Nature methods 9, 245–253 (2012).

59. Woodier J, Rainbow RD, Stewart AJ, Pitt SJ. Intracellular zinc modulates cardiac ryanodine receptor-mediated calcium release. Journal of Biological Chemistry 290, 17599–17610 (2015).

60. DeJesus MA, Ambadipudi C, Baker R, Sassetti C, Ioerger TR. TRANSIT--A Software Tool for Himar1 TnSeq Analysis. PLoS Comput Biol 11, e1004401 (2015).

61. Liao Y, Smyth GK, Shi W. featureCounts: an efficient general purpose program for assigning sequence reads to genomic features. Bioinformatics 30, 923–930 (2014).

62. Robinson MD, McCarthy DJ, Smyth GK. edgeR: a Bioconductor package for differential expression analysis of digital gene expression data. Bioinformatics 26, 139–140 (2010).

63. Nugent T, Jones DT. Transmembrane protein topology prediction using support vector machines. BMC Bioinformatics 10, 159 (2009).

64. Lomize MA, Pogozheva ID, Joo H, Mosberg HI, Lomize AL. OPM database and PPM web server: resources for positioning of proteins in membranes. Nucleic Acids Res 40, D370–D376 (2012).

65. Jo S, Kim T, Iyer VG, Im W. CHARMM-GUI: a web-based graphical user interface for CHARMM. J Comput Chem 29, 1859–1865 (2008).

66. Jorgensen WL, Chandrasekhar J, Madura JD, Impey RW, Klein ML. Comparison of Simple Potential Functions for Simulating Liquid Water. J Chem Phys 79, 926–935 (1983).

67. Huang J, et al. CHARMM36m: an improved force field for folded and intrinsically disordered proteins. Nat Methods 14, 71–73 (2017).

68. Van der Spoel D, Lindahl E, Hess B, Groenhof G, Mark AE, Berendsen HJC. GROMACS: Fast, flexible, and free. Journal of Computational Chemistry 26, 1701–1718 (2005).

69. Parrinello M, Rahman A. Polymorphic Transitions in Single-Crystals - a New Molecular-Dynamics Method. J Appl Phys 52, 7182–7190 (1981).

70. Evans DJ, Holian BL. The Nose-Hoover Thermostat. J Chem Phys 83, 4069–4074 (1985).

71. Darden T, York D, Pedersen L. Particle Mesh Ewald - an N.Log(N) Method for Ewald Sums in Large Systems. J Chem Phys 98, 10089–10092 (1993).

72. Hess B, Bekker H, Berendsen HJC, Fraaije JGEM. LINCS: A linear constraint solver for molecular simulations. Journal of Computational Chemistry 18, 1463–1472 (1997).

73. Aksimentiev A, Schulten K. Imaging alpha-hemolysin with molecular dynamics: ionic conductance, osmotic permeability, and the electrostatic potential map. Biophys J 88, 3745–3761 (2005).

74. UniProt C. UniProt: the Universal Protein Knowledgebase in 2023. Nucleic Acids Res 51, D523–D531 (2023).

75. Eddy SR. Accelerated profile HMM searches. PLoS computational biology 7, e1002195 (2011).

76. Camacho C, et al. BLAST+: architecture and applications. BMC Bioinformatics 10, 421 (2009).

77. Williams DJ. hamburger doi:10.5281/zenodo.6981393. (2022).

78. Abby SS, Cury J, Guglielmini J, Neron B, Touchon M, Rocha EP. Identification of protein secretion systems in bacterial genomes. Scientific reports 6, 23080 (2016).

79. Nguyen L-T, Schmidt HA, von Haeseler A, Minh BQ. IQ-TREE: a fast and effective stochastic algorithm for estimating maximum-likelihood phylogenies. Molecular biology and evolution 32, 268–274 (2014).

80. Hoang DT, Chernomor O, von Haeseler A, Minh BQ, Vinh LS. UFBoot2: Improving the Ultrafast Bootstrap Approximation. Mol Biol Evol 35, 518–522 (2018).

81. Kalyaanamoorthy S, Minh BQ, Wong TKF, von Haeseler A, Jermiin LS. ModelFinder: fast model selection for accurate phylogenetic estimates. Nat Methods 14, 587–589 (2017).

82. Yu G, Smith DK, Zhu H, Guan Y, Lam TTY. ggtree: an R package for visualization and annotation of phylogenetic trees with their covariates and other associated data. Methods in Ecology and Evolution 8, 28–36 (2017).

83. Wickham H. ggplot2: elegant graphics for data analysis. Springer (2016).

