## Supplementary Data for "A widely-occurring family of pore-forming effectors broadens the impact of the *Serratia* Type VI secretion system"

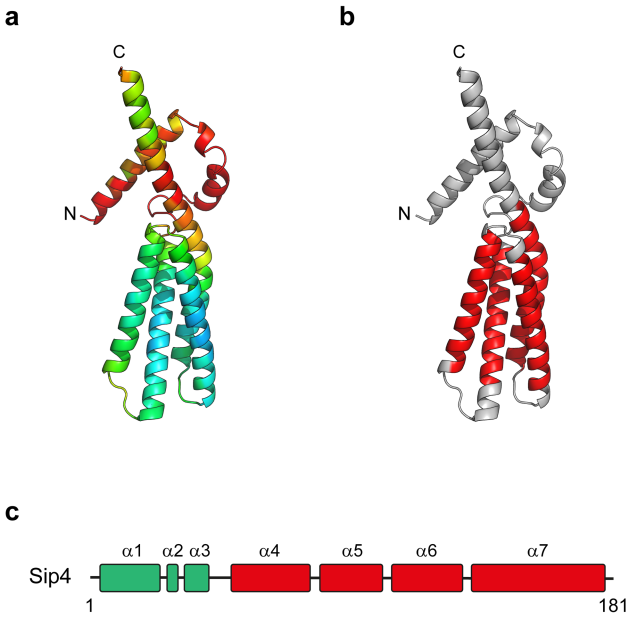


**Supplementary Figure 1. The structure of Sip4 predicted by AlphaFold2**. (a) Structure coloured by pLDDT value, spectrum from red (<50), yellow, green, cyan, to blue (>90). (b) Regions predicted to form transmembrane helices by MEMSAT2 are highlighted in red (amino acids 53-74 in α4, amino acids 87-107 in α5, amino acids 114-135 in α6, and amino acids142-168 in α7). (c) Secondary structure elements in the predicted structure of Sip4, with α-helical regions predicted to include transmembrane helices coloured red.

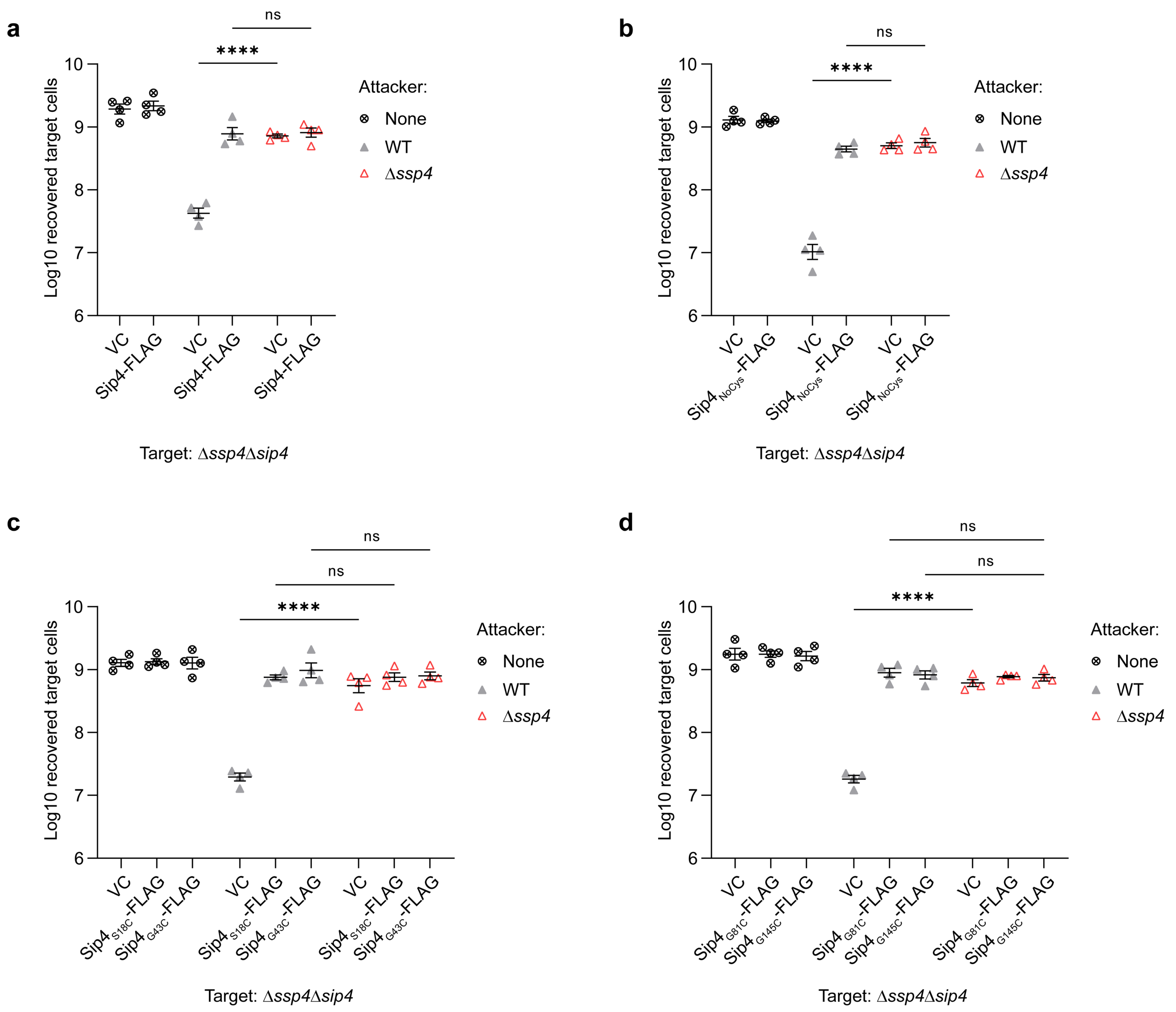


**Supplementary Figure 2. Removal of native cysteine residues and substitution of selected other residues with cysteine for mPEG-Mal topology mapping does not affect Sip4 functionality.** Recovery of *S. marcescens* Db10 Δ*ssp4* Δ*sip4* target strains carrying the vector control (VC, pSUPROM) or plasmids directing the expression of (a) wild type Sip4 with C-terminal 3xFLAG tag (Sip4-FLAG), (b) a derivative lacking native Cys residues (Sip4$_{NoCys}$-FLAG, C60A-C127A-C128A), (c) derivatives of Sip4$_{NoCys}$-FLAG with S18C or G43C substitutions (Sip4$_{S18C}$-FLAG, Sip4$_{G43C}$-FLAG), or (d) derivatives of Sip4$_{NoCys}$-FLAG with G81C or G145C substitutions (Sip4$_{G81C}$-FLAG, Sip4$_{G145C}$-FLAG), following co-culture with wild type (WT) or Δ*ssp4* attacker strains of Db10. None, no attacker. Data are presented as mean +/- SEM with individual data points overlaid (n=4 biological replicates; **** $p<0.0001$, ns not significant, one-way ANOVA with Tukey's test; for clarity, only selected comparisons are displayed).

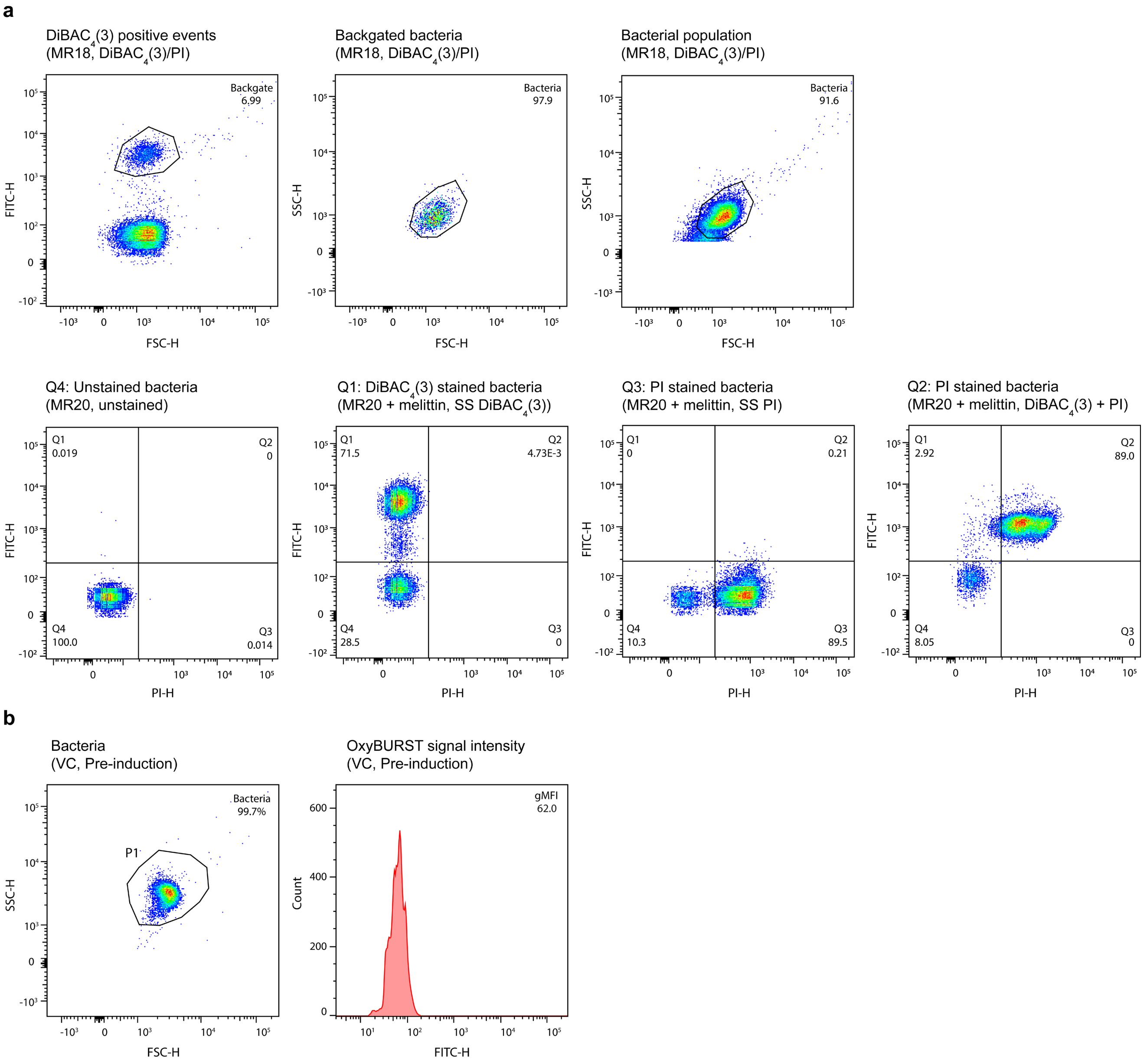


**Supplementary Figure 3. Illustration of the gating strategy used in flow cytometry experiments.** (a) Depolarisation/permeabilisation assays. The bacterial population was identified from the melittin treated, singly stained (SS) DiBAC$_4$(3) samples and used to determine the forward scatter (FSC) / side scatter (SSC) profile of the cognate population in all cases. (b) OxyBURST Green staining assays. The bacterial population was identified using FSC / SSC. In both panels, the sample and staining protocol used to define each population is provided in brackets.

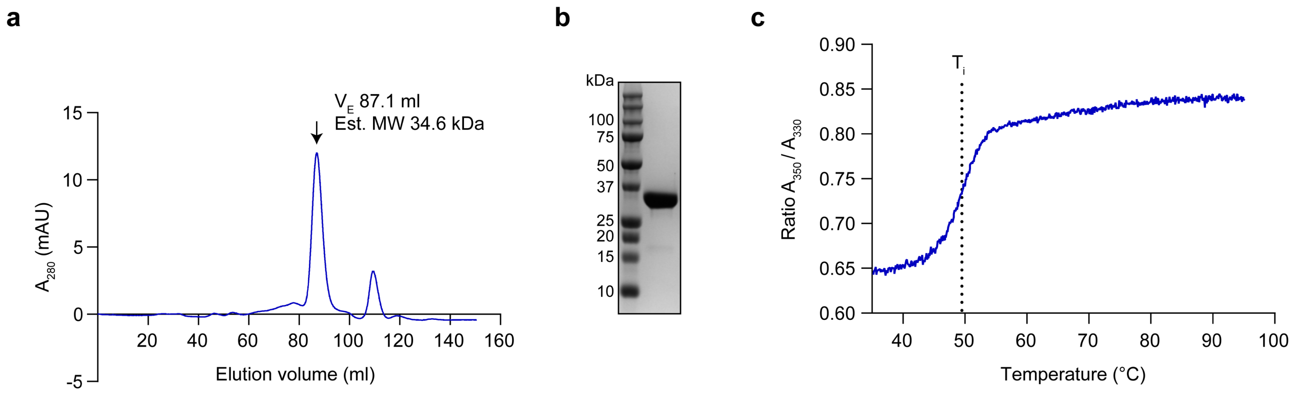


**Supplementary Figure 4. Purification and analysis of Ssp4.** Overproduced $His_6$-GST-Ssp4 protein was purified to apparent homogeneity by Ni-NTA and GST-affinity chromatography. Following cleavage of His-GST, purified Ssp4 was separated by size exclusion chromatography (SEC). (a) SEC profile of purified Ssp4 separated using a Superdex 200 Hiload 26/600 column. The elution volume ($V_E$) and estimated molecular weight (Est. Mw) of the Ssp4 peak, based on calibration of the column with protein standards, is indicated. (b) Protein within the Ssp4 peak was visualised by SDS-PAGE and Coomassie staining. The purified Ssp4 protein has a predicted Mw of 35.5 kDa. (c) The unfolding profile of purified Ssp4 was determined by thermal calorimetry. The inflection temperature (Ti, 49.5 °C) is displayed as a dotted line.

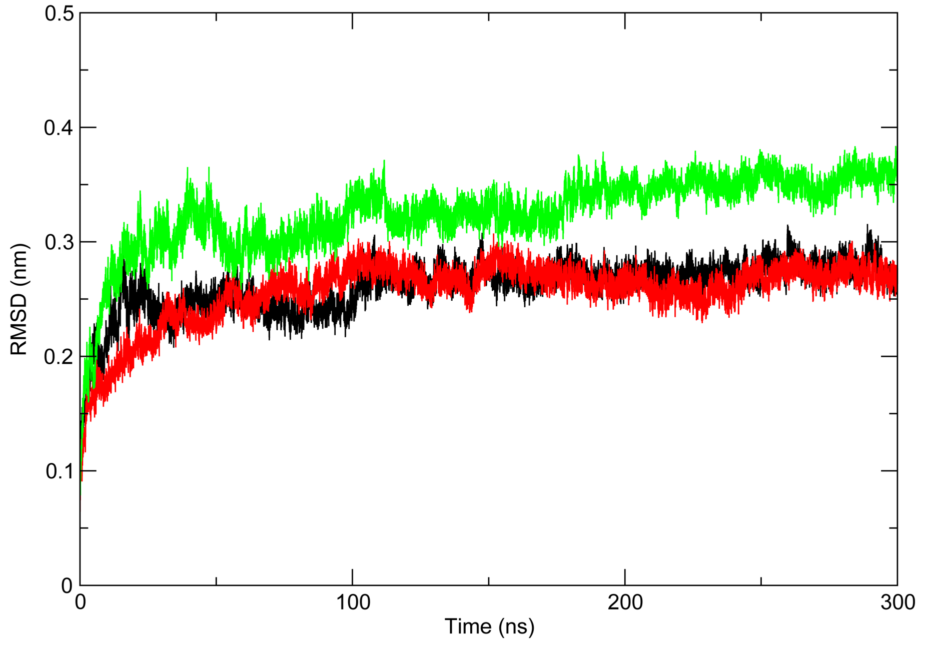


**Supplementary Figure 5. Structural stability of the Ssp4$_{114\text{-}302}$ tetramer in a model lipid bilayer.** Root-mean-square deviation (C-alpha after least squares fit to C-alpha) over a simulation time of 300 ns for the Ssp4$_{114\text{-}302}$ tetramer embedded in a POPC bilayer in three independent simulations.

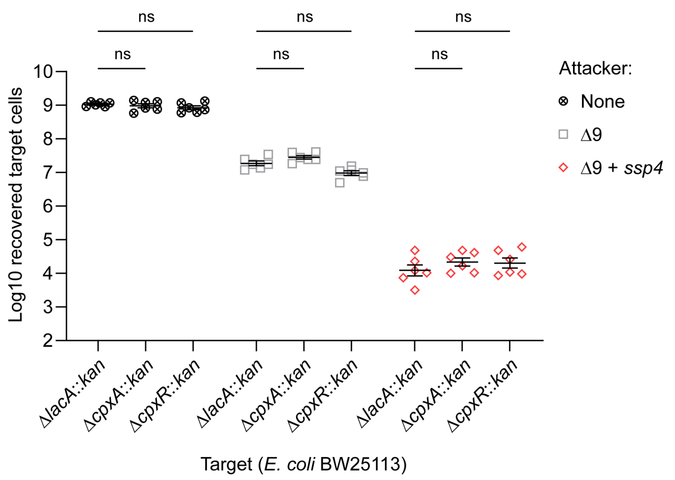


**Supplementary Figure 6. Deletion of *cpxA* or *cpxR* in *E. coli* has no detectable impact on susceptibility to Ssp4 intoxication.** Recovery of *E. coli* BW25112 carrying control (Δ*lacA*::*kan*) or Cpx (Δ*cpxA*::*kan*, Δ*cpxR*::*kan*) gene deletions, following co-culture with attacking strains of *S. marcescens* Db10 lacking known antibacterial effectors (Δ9) or delivering only Ssp4 (Δ9 + *ssp4*). None, no attacker; *kan*, kanamycin-resistance gene replacing deleted gene and providing selection for target cells. Data are presented as mean ± SEM with individual data points overlaid (n=6 biological replicates; ns, not significant; one-way ANOVA with Tukey's test; for clarity, only selected comparisons are displayed).

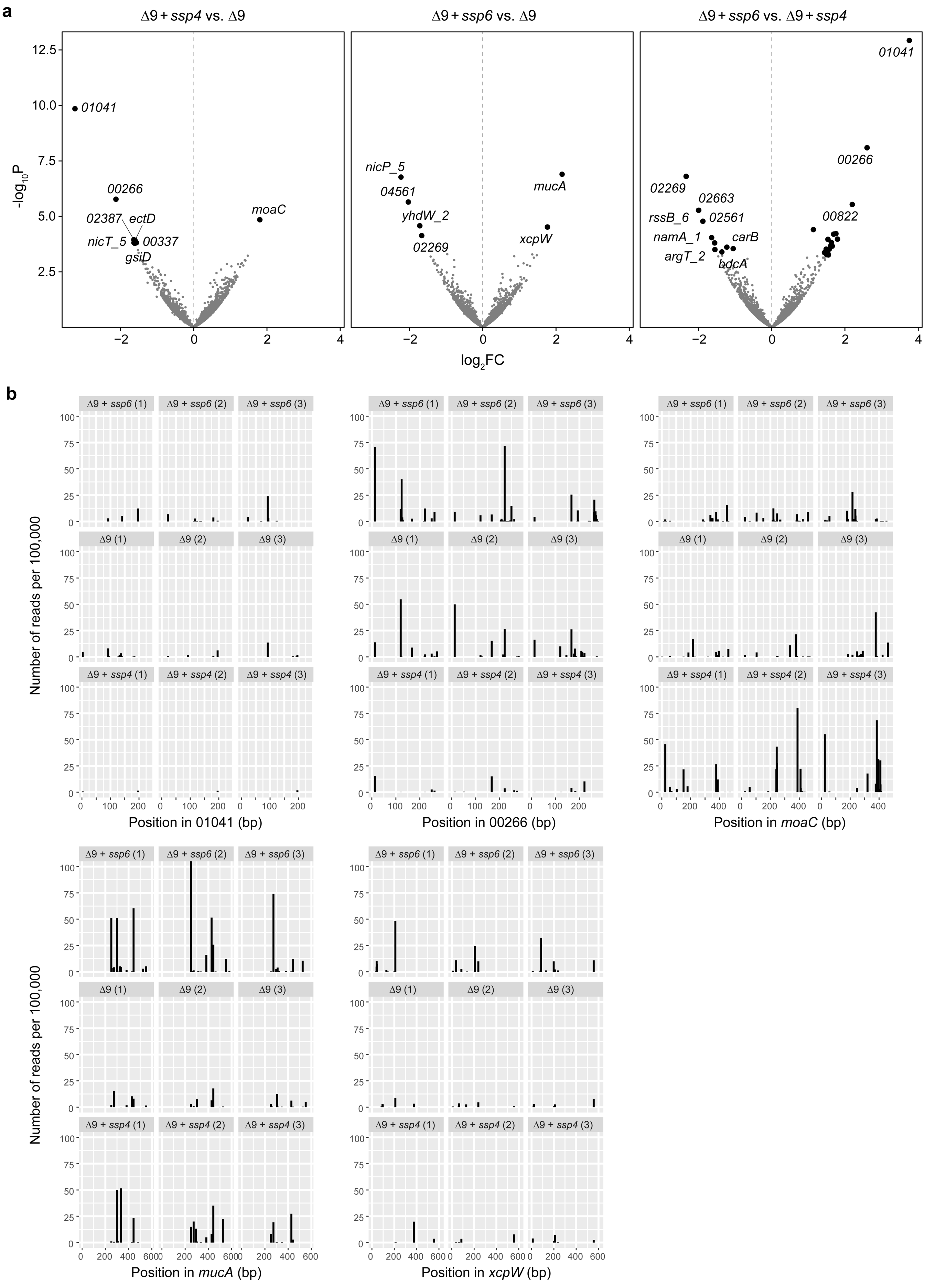
a
Δ9 + ssp4 vs. Δ9
Δ9 + ssp6 vs. Δ9
Δ9 + ssp6 vs. Δ9 + ssp4
-log10P
log2FC
01041
00266
02387
ectD
nicT_5
00337
gsiD
moaC
nicP_5
04561
yhdW_2
02269
mucA
xcpW
02269
02663
rssB_6
02561
namA_1
carB
argT_2
bdcA
00266
00822
01041
b
Number of reads per 100,000
Δ9 + ssp6 (1)
Δ9 + ssp6 (2)
Δ9 + ssp6 (3)
Δ9 (1)
Δ9 (2)
Δ9 (3)
Δ9 + ssp4 (1)
Δ9 + ssp4 (2)
Δ9 + ssp4 (3)
Position in 01041 (bp)
Position in 00266 (bp)
Position in moaC (bp)
Position in mucA (bp)
Position in xcpW (bp)

**Supplementary Figure 7. All three Tn-seq pairwise comparisons and individual insertion sites in genes of interest.** (a) Volcano plots summarising the change in recovery of *P. fluorescens* 55 transposon insertion mutants between control (Δ9) and Ssp4-delivering attackers (left), between control and Ssp6-delivering attackers (middle), and between Ssp4- and Ssp6-delivering attackers (right) on a per gene basis. Log2 fold change in normalized read count is plotted against -log10 p-value and genes significantly altered between condition (FDR <0.05) are highlighted as black dots, with all (left, middle) or selected (right) gene annotations. (b) Position and number of sequencing reads for individual transposon insertion sites across five genes of interest in each replicate for each attacking strain.

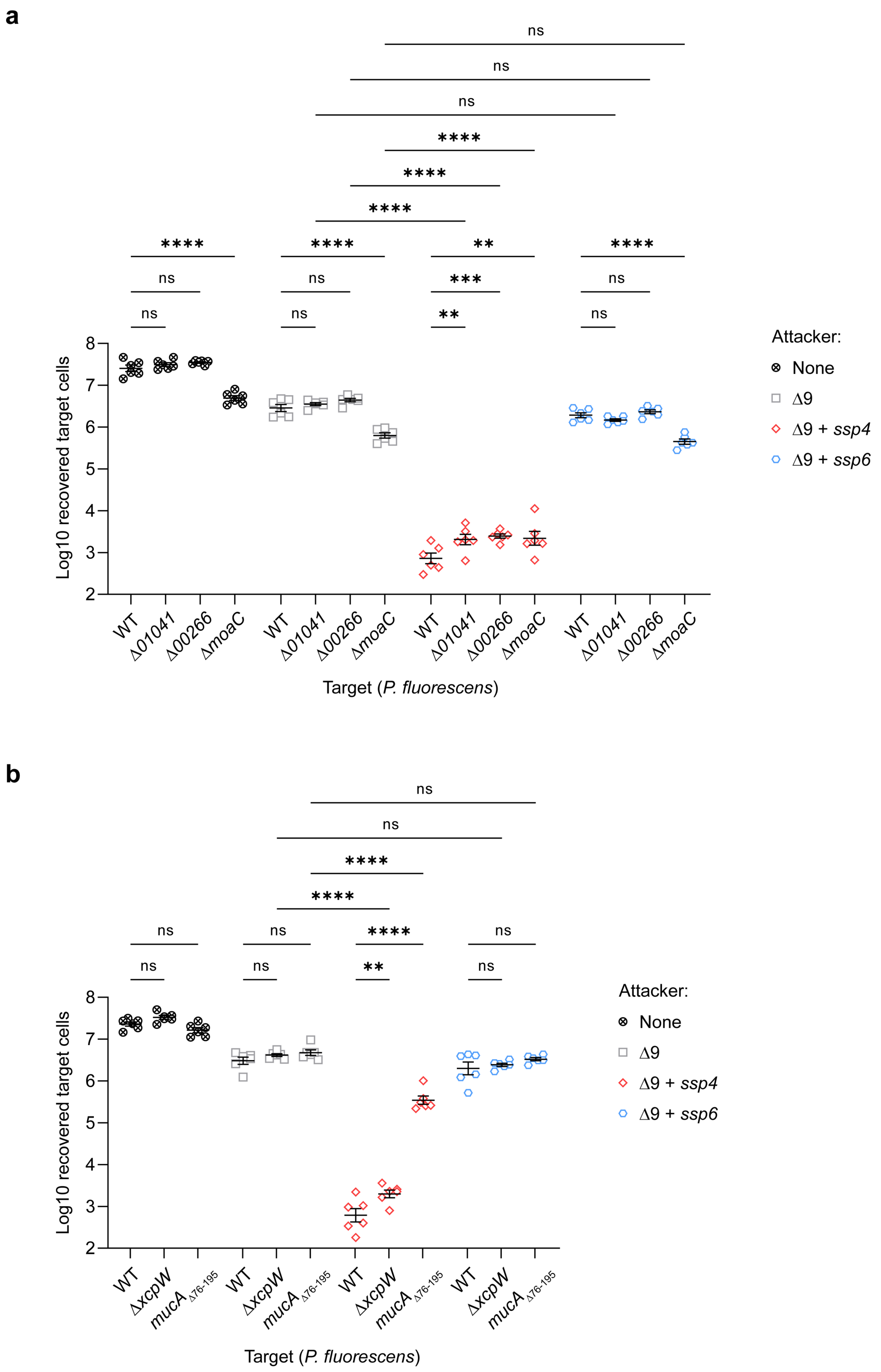


**Supplementary Figure 8. Validation of mutations identified by Tn-seq in *P. fluorescens* as potentially affecting susceptibility to Ssp4 or Ssp6.** Recovery of wild type (WT) or reconstructed defined mutants (*Δ01041, Δ00266, ΔmoaC*, panel a, or *ΔxcpW, mucA*$_{\Delta76\text{-}195}$, panel b) of *P. fluorescens* 55, following co-culture with attacking strains of *S. marcescens* Db10 as indicated. None, no attacker. Data are presented as mean ± SEM with individual data points overlaid (n=6 biological replicates; **** P<0.0001, *** P<0.001, ** P<0.01, ns, not significant; one-way ANOVA with Tukey's test; for clarity, only selected comparisons are displayed). Panels a and b represent the full experiments from which Figure 7c and 7d, respectively, are taken.

**Supplementary Table 1. Strains and plasmids used in this study.**

| Name | Description | Reference / Source |
|---|---|---|
| **Bacterial strains** | | |
| ***Serratia marcescens*** | | |
| Db10 | Wild type strain | [1] |
| SJC11 | Db10 Δ*tssE* (Δ*SMDB11_2271*) | [2] |
| MJF8 | Db10 Δ*ssp4* (Δ*SMDB11_3980*) | [3] |
| JAD01 | Db10 Δ*ssp4*Δ*sip4* (Δ*SMDB11_3980-3979*) | [3] |
| JAD06/AO01 | Db10 Δ*ssp4*Δ*sip4* (Δ*SMDB11_3980-3979*), $Str^R$ | [3] |
| AO03 | Db10 Δ*ssp4*Δ*sip4* Δ*lacZ*::$P_{T5}$-*gfpmut2* $kan^R$. Encodes cytoplasmic GFP, IPTG-inducible and expressed constitutively at a low level. | This study |
| AO07 | Db10 Δ*lacZ*::$P_{T5}$-*mCherry* $kan^R$. Encodes cytoplasmic mCherry, IPTG-inducible and expressed constitutively at a low level. | [4] |
| AO08 | Db10 Δ*tssE* Δ*lacZ*::$P_{T5}$-*mCherry* $kan^R$ | [4] |
| AO09 | Db10 Δ*ssp4* Δ*lacZ*::$P_{T5}$-*mCherry* $kan^R$ | This study |
| YL37 | Db10 Δ9 [Δ*ssp1* (Δ*SMDB11_2261*), Δ*ssp2* (Δ*SMDB11_2264*), Δ*ssp3*/*tfe1* (Δ*SMDB11_1112*), Δ*ssp4* (Δ*SMDB11_3980*), Δ*ssp5* (Δ*SMDB11_4628*), Δ*ssp6* (Δ*SMDB11_4673*), Δ*rhs1* (Δ*SMDB11_2278*), *rhs2*$_{H1369A}$ (*SMDB11_1610*$_{H1369A}$), Δ*slp* (Δ*SMDB11_0927*)] | This study; constituent mutations[5, 6, 7] |
| YL57 | Db10 Δ9 + *ssp4* (Δ*ssp1* Δ*ssp2* Δ*ssp3*/*tfe* Δ*ssp5* Δ*ssp6* Δ*rhs1* *rhs2*$_{H1369A}$ Δ*slp*); generated by restoring the wild type *ssp4* allele in YL37 | This study |
| GM103 | Db10 Δ9 + *ssp6* (Δ*ssp1* Δ*ssp2* Δ*ssp3*/*tfe1* Δ*ssp4* Δ*ssp5* Δ*rhs1* *rhs2*$_{H1369A}$ Δ*slp*); generated by restoring the wild type *ssp6* allele in YL37 | This study |
| YL56 | Db10 Δ9 + *ssp2* (Δ*ssp1* Δ*ssp3*/*tfe1* Δ*ssp4* Δ*ssp5* Δ*ssp6* Δ*rhs1* *rhs2*$_{H1369A}$ Δ*slp*); generated by restoring the wild type *ssp2* allele in YL37 | This study |
| YL33 | Db10 Δ9 + *rhs2* (Δ*ssp1* Δ*ssp2* Δ*ssp3*/*tfe1* Δ*ssp4* Δ*ssp5* Δ*ssp6* Δ*rhs1* Δ*slp*) | This study; constituent mutations[5, 6, 7] |
| ***Pseudomonas fluorescens*** | | |
| *P. fluorescens* 55 | Wild type strain | [2] |
| 55 (mScarlet) | *P. fluorescens 55* $P_{rpsG}$-mScarlet $Gen^R$ | This study |
| MR13 (mScarlet) | *P. fluorescens 55* Δ*01041* (Δ*33931E_Pfluorescens55_01041*), $P_{rpsG}$-mScarlet $Gen^R$ | This study |
| MR14 (mScarlet) | *P. fluorescens 55* Δ*00266* (Δ*33931E_Pfluorescens55_00266*), $P_{rpsG}$-mScarlet $Gen^R$ | This study |
| MR15 (mScarlet) | *P. fluorescens 55* Δ*moaC* (Δ*33931E_Pfluorescens55_01260*), $P_{rpsG}$-mScarlet $Gen^R$ | This study |
| MR16 (mScarlet) | *P. fluorescens 55* Δ*xcpW* (Δ*33931E_Pfluorescens55_02495*), $P_{rpsG}$-mScarlet $Gen^R$ | This study |
| MR17 (mScarlet) | *P. fluorescens 55* *mucA*$_{\Delta 76\text{-}195}$ (*33931E_Pfluorescens55_01704*$_{\Delta 76\text{-}195}$), encodes MucA lacking C-terminal 120 amino acids; $P_{rpsG}$-mScarlet $Gen^R$ | This study |
| MR18 | *P. fluorescens 55* *att*Tn*7*::$P_{RhaBAD}$-*sp-ssp4*, $Gen^R$ | This study |
| MR19 | *P. fluorescens 55* *att*Tn*7*::$P_{RhaBAD}$-*sp-ssp6*, $Gen^R$ | This study |
| MR20 | *P. fluorescens 55* *att*Tn*7*::$P_{RhaBAD}$, $Gen^R$ | This study |
| ***Escherichia coli*** | | |
| MG1655 | Wild type (model K-12 strain) | [8] |
| CC118λ*pir* | Donor strain for pKNG101-derived allelic exchange plasmids (λ*pir*) | [9] |
| HH26 pNJ5000 | Helper strain for mobilization of pKNG101 | [10] |
| SM10λpir | Donor strain for conjugal transfer of pMQ30, pTNS1 and pIT2 | [11] |
| HB101 pRK2013 | Helper strain for mobilization of pUC18T-miniTn7T-$Gen^R$ and pJM220 | [11] |

| | | |
|---|---|---|
| BL21(DE3) | Protein overexpression strain. Chromosomal λDE3 encodes IPTG-inducible T7 RNA polymerase. | Novagen |
| sHuffle T7 | *E. coli* K-12 expressing DsbC to promote disulfide bond formation. | NEB |
| BW25113 | Parental strain of the Keio collection. Δ(*araD-araB*)*567*, Δ*lacZ4787 (::rrnB-4), lacIp-4000(lacI*$^{Q}$*), λ*$^{-}$*, rpoS369(Am), rph-1,* Δ*(rhaD-rhaB)568, hsdR514* | [12] |
| - | BW25113 Δ*lacA*::Kan$^{R}$ (*b0342*) from Keio collection | [12] |
| - | BW25113 Δ*cpxA*::Kan$^{R}$ (*b3911*) from Keio collection | [12] |
| - | BW25113 Δ*cpxR*::Kan$^{R}$ (*b3912*) from Keio collection | [12] |
| MR21 | BW25113 Δ*lacA* | This study |
| MR22 | BW25113 Δ*cpxA* | This study |
| MR23 | BW25113 Δ*cpxR* | This study |
| **<u>Plasmids</u>** | | |
| **Mutant generation** | | |
| pKNG101 | Suicide vector for allelic exchange in *S. marcescens* (Str$^{R}$, *sacBR, mobRK2, oriR6K*) | [13] |
| pSAN72 | pKNG101-derived allelic exchange plasmid for the generation of chromosomal Δ*lacZ*::$P_{T5}$-*gfpmut2 kan*$^{R}$ | [14] |
| pSC1706 | pKNG101-derived allelic exchange plasmid for the generation of chromosomal Δ*lacZ*::$P_{T5}$-*mCherry kan*$^{R}$ | [4] |
| pSC1829 | pKNG101-derived allelic exchange plasmid for restoration of the wild type *ssp2* allele in YL37 | This study |
| pSC1830 | pKNG101-derived allelic exchange plasmid for restoration of the wild type *ssp4* allele in YL37 | This study |
| pSC2526 | pKNG101-derived allelic exchange plasmid for restoration of the wild type *ssp6* allele in YL37 | This study |
| pMQ30 | Suicide vector for allelic exchange in *P. fluorescens* (Amp$^{R}$, *sacBR, mobIncP, oriColE1*) | [15] |
| pSC3443 | pMQ30-derived allelic exchange plasmid for the generation of chromosomal in-frame Δ*33931E_Pfluorescens55_01041* deletion | This study |
| pSC3444 | pMQ30-derived allelic exchange plasmid for the generation of chromosomal in-frame Δ*33931E_Pfluorescens55_00266* deletion | This study |
| pSC3445 | pMQ30-derived allelic exchange plasmid for the generation of chromosomal in-frame Δ*33931E_Pfluorescens55_01260* | This study |
| pSC3447 | pMQ30-derived allelic exchange plasmid for the generation of chromosomal in-frame Δ*33931E_Pfluorescens55_02495* deletion | This study |
| pSC3454 | pMQ30-derived allelic exchange plasmid for the generation of chromosomal in-frame deletion of amino acids 76-195 in MucA (33931E_Pfluorescens55_01704) | This study |
| pUC18T-miniTn*7*T-Gm$^{R}$ $P_{rpsG}$-mScarlet | Suicide vector for integration of gentamycin resistance cassette and constitutively expressed mScarlet into *att*Tn*7* site of *P. fluorescens* (Gen$^{R}$, Amp$^{R}$, *oriPMB1*) | [16] |
| pTNS1 | Plasmid directing Tn*7* transposase expression for Tn*7* integration in *P. fluorescens* | [17] |
| pJM220 | Suicide vector for integration of gentamycin resistance cassette and gene of interest under the control of rhamnose-inducible promoter in the *att*Tn*7* site of *P. fluorescens* (Gen$^{R}$, Amp$^{R}$, *oriPMB1*, *rhaSR*-$P_{RhaBAD}$). | [18] |
| pSC3457 | Coding sequence for a fusion of the N-terminal signal peptide from *E. coli* OmpA to Ssp4 (sp-Ssp4) in pJM220 | This study |
| pSC3459 | Coding sequence for a fusion of the N-terminal signal peptide from *E. coli* OmpA to Ssp6 (sp-Ssp4) in pJM220 | This study |
| pCP20 | Temperature-sensitive plasmid for thermal induction of FLP recombinase (Amp$^{R}$, Cml$^{R}$, *ori pSC101*) | [19] |
| pIT2 | Vector containing Tn*5*-based transposon T8 (IS*lacZ*/hah-*tet*) for generation of transposon library in *P. fluorescens* | [20] |

| | | |
|---|---|---|
| **Heterologous gene expression in *E. coli*** | | |
| pBAD18-Kn | Arabinose-inducible expression vector; gene of interest is cloned downstream of the $P_{ara}$ promoter ($Kan^R$) | [21] |
| pSC1234 | Coding sequence for a fusion of the N-terminal signal peptide from *E. coli* OmpA to Ssp4 (sp-Ssp4) in pBAD18-Kn | [3] |
| pSC861 | Coding sequence for sp-Ssp4 + Sip4 in pBAD18-Kn | [3] |
| pSC1236 | Coding sequence for a fusion of the N-terminal signal peptide from *E. coli* OmpA to Ssp6 (sp-Ssp6) in in pBAD18-Kn | [3] |
| pSC1271 | Coding sequence for sp-Ssp6 + Sip6 in pBAD18-Kn | [4] |
| pSC838 | Coding sequence for Ssp5 in pBAD18-Kn | [3] |
| pSC839 | Coding sequence for Ssp5 + Sip5a in pBAD18-Kn | [3] |
| **Protein production** | | |
| pSC3801 | Vector for protein overproduction under the control of the T7 promoter. Permits fusion of an $His_{10}$ tag followed by a TEV protease cleavage site to the N-terminus of the overproduced protein ($Amp^R$). Derived from pET15b-TEV[22] with $His_{10}$ replacing $His_6$. | This study, [22] |
| pSC3455 | Coding sequence for Ssp4 in pSC3801, for production of TEV-cleavable N-terminal $His_{10}$-tagged Ssp4 | This study |
| pSC3456 | Coding sequence for Ssp6 in pSC3801, for production of TEV-cleavable N-terminal $His_{10}$-tagged Ssp6 | This study |
| pHis-GEX-6P-1 | Protein overexpression vector for fusion with PreScission-cleavable N-terminal $His_6$/GST tag under the control of the Tac promoter. Derived from pGEX-6P-1 ($Amp^R$) | van Aalten lab |
| pSC3407 | Coding sequence for Ssp4 in pHis-GEX-6P-1, for production of PreScission-cleavable N-terminal $His_6$-GST-tagged Ssp4 | This study |
| pSC3460 | Coding sequence for Ssp6 in pHis-GEX-6P-1, for production of PreScission-cleavable N-terminal $His_6$-GST-tagged Ssp6 | This study |
| **Gene expression *in trans* in *S. marcescens*** | | |
| pSUPROM | Vector for constitutive expression of cloned genes under the control of the *E. coli tat* promoter ($Kan^R$) | [23] |
| pSC2305 | Coding sequence for Sip4 with a C-terminal 3xFLAG tag (Sip4-FLAG) in pSUPROM | This study |
| pSC2310 | Coding sequence for Sip4 with a C-terminal 3xFLAG tag and containing C60A, C127A and C128A substitutions ($Sip4_{NoCys}$-FLAG) in pSUPROM | This study |
| pSC2315 | pSC2310 with S18C substitution in $Sip4_{NoCys}$-FLAG | This study |
| pSC2316 | pSC2310 with G43C substitution in $Sip4_{NoCys}$-FLAG | This study |
| pSC2320 | pSC2310 with G81C substitution in $Sip4_{NoCys}$-FLAG | This study |
| pSC2321 | pSC2310 with G145C substitution in $Sip4_{NoCys}$-FLAG | This study |

### Supplementary Table 2. Oligonucleotide primers used in this study.

| Plasmid | Sequence of relevant primers (5'-3') | Description |
|---|---|---|
| pSC1829 | TATATCTAGACCACACTTGCAATTCCTGC | Forward primer to clone *ssp2* and flanking regions into pKNG101 |
| | TATAGGGCCCTTGTTCAAATGCATCCATCG | Reverse primer to clone *ssp2* and flanking regions into pKNG101 |
| pSC1830 | TATATCTAGAAGCTTCCTCAAGTTCTGCCAAC | Forward primer to clone *ssp4* and flanking regions into pKNG101 |
| | TATAGGGCCCAAGTTTTTCCGTTCTGACGGTTTATC | Reverse primer to clone *ssp4* and flanking regions into pKNG101 |
| pSC2526 | TATATCTAGAGGGATCAGTTCGATGTGCG | Forward primer to clone *ssp6* and flanking regions into pKNG101 |
| | TATAGGGCCCCGACGACATCAAGTACCTGTTCG | Reverse primer to clone *ssp6* and flanking regions into pKNG101 |
| pSC3443 | ATATCTAGAGAATGTGGAGTGGTGGCAC | Upstream forward primer for 01041 deletion in *P. fluorescens 55* |
| | ATAGGATCCCATCAGATACTCTCCCGTTC | Upstream reverse primer for 01041 deletion in *P. fluorescens 55* |
| | ATAGGATCCGAGCCAGCTTCGAATTGACAT | Downstream forward primer for 01041 deletion in *P. fluorescens 55* |
| | ATAGAATTCCGGGCAAGTCACGTATGTG | Downstream reverse primer for 01041 deletion in *P. fluorescens 55* |
| pSC3444 | ATATCTAGACCTGGATTCGCATCTGCG | Upstream forward primer for 00266 deletion in *P. fluorescens 55* |
| | ATAGTCGACAGTCATGCTGCTGTTCTCC | Upstream reverse primer for 00266 deletion in *P. fluorescens 55* |
| | ATAGTCGACGTGTTCTAGGCACCAAATGG | Downstream forward primer for 00266 deletion in *P. fluorescens 55* |
| | ATAGAATTCTAATCGGCATGCCGCGTC | Downstream reverse primer for 00266 deletion in *P. fluorescens 55* |
| pSC3445 | ATATCTAGACAGGAAATCGGCTTCCTGC | Upstream forward primer for *moaC* deletion in *P. fluorescens 55* |
| | ATAGGATCCATCGAGATGAGTCAGCACG | Upstream reverse primer for *moaC* deletion in *P. fluorescens 55* |
| | ATAGGATCCGCATGAGCATCAACGTATTGT | Downstream forward primer for *moaC* deletion in *P. fluorescens 55* |
| | ATAGAATTCCGCGTCTTCAAGTAATCCATC | Downstream reverse primer for *moaC* deletion in *P. fluorescens 55* |
| pSC3447 | ATATCTAGAGGTGAAACGATGGGCGAATG | Upstream forward primer for *xcpW* deletion in *P. fluorescens 55* |
| | ATAGGATCCCTGATTCATGGCAGGCGC | Upstream reverse primer for *xcpW* deletion in *P. fluorescens 55* |
| | ATAGGATCCCCGTTGAACTAACTCAACAACAC | Downstream forward primer for *xcpW* deletion in *P. fluorescens 55* |
| | ATAGAATTCGAGCAACGCCAAGTGGTG | Downstream reverse primer for *xcpW* deletion in *P. fluorescens 55* |
| pSC3454 | ATATCTAGAGGTGATAGCGCGTTTTACAC | Upstream forward primer for deletion of part of *mucA* gene encoding MucA amino acids 76–195 in *P. fluorescens 55* |
| | ATAGTCGACGGCATCACGACTCATGGC | Upstream reverse primer for deletion of part of *mucA* gene encoding MucA amino acids 76–195 in *P. fluorescens 55* |
| | ATAGTCGACTGCCGGAACGGCTTCATC | Downstream forward primer for deletion of part of *mucA* gene encoding MucA amino acids 76–195 in *P. fluorescens 55* |
| | ATAGAATTCTGCAGATCGCGATCATCCG | Downstream reverse primer for deletion of part of *mucA* gene encoding MucA amino acids 76–195 in *P. fluorescens 55* |
| pSC3457 | TTTACTAGTGCTAGCGAATTCGAGCTC | Forward primer to amplify sequence encoding sp-Ssp4 for cloning into pJM220 |
| | TTTGGGCCCTACCTCACCATCTGCGG | Reverse primer to amplify sequence encoding sp-Ssp4 for cloning into pJM220 (final insert sequence below) |
| pSC3459 | TTTACTAGTGCTAGCGAATTCGAGCTC | Forward primer to amplify sequence encoding sp-Ssp6 for cloning into pJM220 |
| | TTTGGGCCCTTTCGACACCTTTTCAAAAAATAG | Reverse primer to amplify sequence encoding sp-Ssp6 from for cloning into pJM220 |

| | | |
|---|---|---|
| | ATAGCCCGGTACCTCGCGAAGGCCT | Forward primer for site-directed mutagenesis for incorporation of stop codon at the end of sp-Ssp6 |
| | TTCGACACCTTTTCAAAAAATAGCTCGGTAATGCG | Reverse primer for site-directed mutagenesis for incorporation of stop codon at the end of sp-Ssp6 (final insert sequence below) |
| pSC3455 | CATATGCTCGAGGATCCG | Universal forward primer for *ssp4 / ssp6* cloning into pSC3801 by Gibson assembly |
| | GCCCTGAAAATACAGGTTTTC | Universal reverse primer for *ssp4 / ssp6* cloning into pSC3801 by Gibson assembly |
| | AAAACCTGTATTTTCAGGGCATGAAAACTGCTTTTTCTACGC | Forward primer for *ssp4* cloning into pSC3801 by Gibson assembly |
| | GCCGGATCCTCGAGCATATGTTACCTCACCATCTGCGG | Reverse primer for *ssp4* cloning into pSC3801 by Gibson assembly |
| pSC3456 | CATATGCTCGAGGATCCG | Universal forward primer for *ssp4 / ssp6* cloning into pSC3801 by Gibson assembly |
| | GCCCTGAAAATACAGGTTTTC | Universal forward primer for *ssp4 / ssp6* cloning into pSC3801 by Gibson assembly |
| | AAAACCTGTATTTTCAGGGCATGGCAAAAGGTGCGAAG | Forward primer for *ssp6* cloning into pSC3801 by Gibson assembly |
| | GCCGGATCCTCGAGCATATGTCATTTCGACACCTTTTCAAAAAATAG | Reverse primer for *ssp6* cloning into pSC3801 by Gibson assembly |
| pSC3407 | GCATGGATCCATGAAAACTGCTTTTTCTACGCC | Forward primer to amplify the sequence encoding Ssp4 for cloning into pHis-GEX-6P-1 |
| | GCATGTCGACTTACCTCACCATCTGCGG | Reverse primer to amplify the sequence encoding Ssp4 for cloning into pHis-GEX-6P-1 |
| pSC3460 | TCTGTTCCAGGGGCCCCTGGCAAAAGGTGCGAAGGAAATC | Forward primer for *ssp6* cloning into BamHI/SalI digested pSC3407 by Gibson assembly |
| | TCACGATGCGGCCGCTCGAGCTATTTCGACACCTTTTCAAAAAATAG | Reverse primer for *ssp6* cloning into BamHI/SalI digested pSC3407 by Gibson assembly |
| pSC2305 | TATAGGATCCGTAATGGCGGAGTGGATGATTAAAC | Forward primer to amplify sequence encoding Sip4-FLAG for cloning into pSUPROM (template sequence below) |
| | TATAGTCGACCGGTTTACTTGTCATCGTCATCC | Reverse primer to amplify sequence encoding Sip4-FLAG for cloning into pSUPROM (template sequence below) |
| pSC2310 | GCATGCTCGGTCTGGCTGCCCTCGGCGCCGC | Forward primer for introduction of C127A and C128A mutations into Sip4-FLAG encoded on pSC2305 by QuikChange site-directed mutagenesis |
| | GCGGCGCCGAGGGCAGCCAGACCGAGCATGC | Reverse primer for introduction of C127A and C128A mutations into Sip4-FLAG encoded on pSC2305 by QuikChange site-directed mutagenesis |
| | TATAGGATCCGTAATGGCGG | Forward primer 1 for introduction of C60A mutation into Sip4-FLAG encoded on pSC2305 by overlap PCR |
| | CGTCAGGGCCACCATC | Reverse primer 1 for introduction of C60A mutation into Sip4-FLAG encoded on pSC2305 by overlap PCR |
| | GATGGTGGCCCTGACG | Forward primer 2 for introduction of C60A mutation into Sip4-FLAG encoded on pSC2305 by overlap PCR |
| | TATAGTCGACCGGTTTACTTGTCATC | Reverse primer 2 for introduction of C60A mutation into Sip4-FLAG encoded on pSC2305 by overlap PCR |
| pSC2315 | TATAGAATTCTGTCGGTTGGCG | Forward primer 1 for introduction of S18C mutation into Sip4-FLAG encoded on pSC2310 by overlap PCR |
| | CGATTTGGCATTCCATCTC | Reverse primer 1 for introduction of S18C mutation into Sip4-FLAG encoded on pSC2310 by overlap PCR |
| | GAGATGGAATGCCAAATCG | Forward primer 2 for introduction of S18C mutation into Sip4-FLAG encoded on pSC2310 by overlap PCR (used with pSC2310 Reverse primer 2) |
| pSC2316 | CGCTGGCACGGCGTC | Reverse primer 1 for introduction of G43C mutation into Sip4-FLAG encoded on pSC2310 by overlap PCR (used with pSC2315 Forward primer 1) |
| | GACGCCGTGCCAGCG | Forward primer 2 for introduction of G43C mutation into Sip4-FLAG encoded on pSC2310 by overlap PCR (used with pSC2310 Reverse primer 2) |
| pSC2320 | GGTCTTGCAGCTCCAGG | Reverse primer 1 for introduction of G81C mutation into Sip4-FLAG encoded on pSC2310 by overlap PCR (used with pSC2315 Forward primer 1) |
| | CCTGGAGCTGCAAGACC | Forward primer 2 for introduction of G81C mutation into Sip4-FLAG encoded on pSC2310 by overlap PCR (used with pSC2310 Reverse primer 2) |

| | | |
|---|---|---|
| pSC2321 | AGGGCGCAGGTGGCG | Reverse primer 1 for introduction of G145C mutation into Sip4-FLAG encoded on pSC2310 by overlap PCR (used with pSC2315 Forward primer 1) |
| | CGCCACCTGCGCCCT | Forward primer 2 for introduction of G145C mutation into Sip4-FLAG encoded on pSC2310 by overlap PCR (used with pSC2310 Reverse primer 2) |

| **Plasmid** | **Template / final insert sequences** |
|---|---|
| pSC2305 | GTAATGGCGGAGTGGATGATTAAACGCCTGATCGTGCAGGGTCGGGAGATGGAAAGCCAAATCGCGGC<br>GCGCGATCGCGCACTTTTCGCCGGTGTGGATGCGCAGCTCGAGCAGCATTTCATGACGCCGGGCCAGCG<br>GTTGTCGCCGCCGGGCGTCGGCAACGTCATGTTGCTGATGGTGTGCCTGACGCTGGGGCTGGCGGGCGT<br>GATGGGTCTGGTCACGGATGTGGCGGCCGCCTGGAGCGGCAAGACCTCCGCCGCCGTGTTGATGGGCAG<br>CGGCGCGATCGTGGCGGTGTGGATGACGTTGATCCTGTTCCAACTGGTGCAGGGTAAAAACAGCGGGGT<br>GGTATTGTTGCAATATTACCTTGGCATGCTCGGTCTGTGTTGCCTCGGCGCCGCGGCGGCGTGGGCGGCG<br>GGCATGACCGGCATCGCCACCGGCGCCCTGCTGCTGGGGGGCGTGGTTGGCGGTGCGCTGTTCAGCAAC<br>CGGGCGGCGTTCTACCTGTATGTCGCCTATTTCCGCACGCGTCGGCGTGTGTTTATCAAACGCCGTTGGC<br>AGAGGGAAGATCTGCGCAACACCCGAGACTACAAAGACCATGACGGTGATTATAAAGATCATGATATC<br>GATTACAAGGATGACGATGACAAGTAAACCG |
| pSC3457 | CTAGTGCTAGCGAATTCGAGCTCAGAGGACGTTAAAATGAAAAAGACAGCTATCGCGATTGCAGTGGCA<br>CTGGCTGGTTTCGCTACCGTAGCGCAGGCCGCTCCGAAATCTAGAAAAACTGCTTTTTCTACGCCGTTCG<br>GTTCTCCCGAAGACGATTTGACCAACGCCGAGTTTGCCCGCCTGTTGGGCAATACCGGCGTACAAGAAT<br>TCTATCGCGATATTAATCTGCAAGCCTTGCAGGACTCGCTGGACTGCAGCCTCGCCTTTCATGGCGTGGG<br>AATTATTTTTTCCGACGGGCAAAACTCATTTTTAGTCCGCCCGACGCAGCACACCGGTTACTCGAACGCT<br>TCGCAGGTTACCCACGTGGTCATCAGCAAAACGGTGGCTCATACCTCGCGCGTTGCCGCGACCAATACG<br>CTGTCGGAAGCGTTGAGCAAACCCAGCGTCAGCAAAGAGCTGGCCTCGGCCGCGCTGTCGTGTGGAACG<br>CTGCTGGTATCGGTCTTTTTACTGGCGTCGGGCAGCGTCGCCGTGCCATTTACCGGCGGTACCAGCTCAG<br>CGGTGGCTTATCTGGGCTATGCCGGCATGGCCGCCAGCGCGTTGCAGTGCGGTAATGGGCTGTACCGCG<br>TCAACAAGCTTTATGACGGGAAAGGCGATGAATTGGCTCAGCTCGATTCAGAGCAGTGGTATATCGCCA<br>CCAGTACCGTGCTCGATGTGATCTCGCTGGCCAGCGCCGGGGCTGCCCTGAAGGAAGCGACGATGACCT<br>ATCGTGCCATGCGCCGCATTTCGGCGCGCAAGGCGACGGAGTGGCTGAAAAGCATGCCCCGCAGCGAA<br>AGGAAGCGACTGACCGAAAACATCATTCGGGCGGAAAATCCGGGAATTTCCAACAACGTTTTGAAAGA<br>GATGGTGAAGAACGGCCTGTACCCGAAACGCTACCCGACGGAAGCGATCCAAAACGGGCTGCGCCAGC<br>AATTGCACTCGGCGCTCAATAACGCGTTGACGTTTGTCGGCAGCGGCATCAGCGGCACCCTGTCGGCGC<br>CGGTGAATGTGAAAACCACCGGGCAGTATTTTGTGGGCATTATGCAGAAATTGCCGCAGATGGTGAGGT<br>AGGGCCC |
| pSC3459 | CTAGTGCTAGCGAATTCGAGCTCAGAGGACGTTAAAATGAAAAAGACAGCTATCGCGATTGCAGTGGCA<br>CTGGCTGGTTTCGCTACCGTAGCGCAGGCCGCTCCGAAATCTAGAGCAAAAGGTGCGAAGGAAATCGCC<br>CAAGAGATGGCGAACGCTGTAAATTCAAAAAGTAATTTCTTTGGCGGTTTTATCGAAGGGGCTATCTCCT<br>TTCCCGTCGATATAGGATACCTGGCATATGATTTTATCAATACAGATAACCGCTCTATAAATCGATATGA<br>CACAGAAAGAATGCTTCGCCTCATTAAAGCTGGACTTGCTAACCAACACTCACTGACCAAAATCGTTAA<br>ACTTGTCGTTGACGAATATCTGAAAAAAGTCGATGTAGATAAAGTCAAAAGATGGGTTGAAAAAGGATC<br>CGGGAAAATTGCAGGTAGGTTTGTGAGCAATCAGGTTCTGATGGTTAACTTAGGGGCGGTGCTTTCCGA<br>ACGGGTGGTTATTCGCCTCGCAACAGGTTATGCCCTAACGTCACTGCTGACCCTCGGAGCTATGAACTCA<br>AGAGCAATCCATACCTCACGCCAGTTACGACAGCGAAACCCTGAAATTTACGATAACTTAAGGCGTGCG<br>GGAAATTTGGATCTCTTATATTTCTGGTAGAACCCAAAACAAAGCCATTTGAACAAGCGATAGAGATTT<br>GGCGAAAAAACAGAGGCGAGTTTGACCGCATTACCGAGCTATTTTTGAAAAGGTGTCGAAATAGCCC |

| **Primer** | **Sequence (5’-3’)** | **Description** |
|---|---|---|
| pIT2 Fwd 1 | CTGGATGGAAAACGGGAAAGGTTCCGTCCA | Forward primer for amplification of C-tailed DNA fragments during Tn-seq |
| olj376 | GTGACTGGAGTTCAGACGTGTGCTCTTCCGATC<br>TGGGGGGGGGGGGGGGG | Reverse primer for amplification of C-tailed DNA fragments during Tn-seq |
| pIT2 Fwd 2 | AATGATACGGCGACCACCGAGATCTACACTCT<br>TTCCCTACACGACGCTCTTCCGATCTGGTTCCG<br>TCCAGGACGCTACTTGTGTATAAGAGT | Forward primer for final amplification of Tn-seq fragments for sequencing. Used with NEBNext index primers |

### References for Supplementary Information


1. Flyg C, Kenne K, Boman HG. Insect pathogenic properties of *Serratia marcescens*: phage-resistant mutants with a decreased resistance to *Cecropia* immunity and a decreased virulence to *Drosophila*. *J Gen Microbiol* **120**, 173-181 (1980).

2. Murdoch SL, Trunk K, English G, Fritsch MJ, Pourkarimi E, Coulthurst SJ. The opportunistic pathogen *Serratia marcescens* utilizes type VI secretion to target bacterial competitors. *J Bacteriol* **193**, 6057-6069 (2011).

3. Fritsch MJ, Trunk K, Diniz JA, Guo M, Trost M, Coulthurst SJ. Proteomic identification of novel secreted antibacterial toxins of the *Serratia marcescens* type VI secretion system. *Mol Cell Proteomics* **12**, 2735-2749 (2013).

4. Mariano G*, et al.* A family of Type VI secretion system effector proteins that form ion-selective pores. *Nature Communications* **10**, 5484 (2019).

5. Alcoforado Diniz J, Coulthurst SJ. Intraspecies Competition in *Serratia marcescens* Is Mediated by Type VI-Secreted Rhs Effectors and a Conserved Effector-Associated Accessory Protein. *J Bacteriol* **197**, 2350-2360 (2015).

6. Cianfanelli FR, Alcoforado Diniz J, Guo M, De Cesare V, Trost M, Coulthurst SJ. VgrG and PAAR Proteins Define Distinct Versions of a Functional Type VI Secretion System. *PLoS Pathog* **12**, e1005735 (2016).

7. Trunk K*, et al.* The type VI secretion system deploys antifungal effectors against microbial competitors. *Nature Microbiology* **3**, 920-931 (2018).

8. Blattner FR*, et al.* The complete genome sequence of *Escherichia coli* K-12. *Science* **277**, 1453-1462 (1997).

9. Herrero M, de Lorenzo V, Timmis KN. Transposon vectors containing non-antibiotic resistance selection markers for cloning and stable chromosomal insertion of foreign genes in gram-negative bacteria. *J Bacteriol* **172**, 6557-6567 (1990).

10. Grinter NJ. A broad-host-range cloning vector transposable to various replicons. *Gene* **21**, 133-143 (1983).

11. Choi KH, Schweizer HP. mini-Tn7 insertion in bacteria with single *att*Tn*7* sites: example *Pseudomonas aeruginosa*. *Nat Protoc* **1**, 153-161 (2006).

12. Baba T*, et al.* Construction of *Escherichia coli* K-12 in-frame, single-gene knockout mutants: the Keio collection. *Mol Syst Biol* **2**, 2006 0008 (2006).

13. Kaniga K, Delor I, Cornelis GR. A wide-host-range suicide vector for improving reverse genetics in Gram-negative bacteria: inactivation of the *blaA* gene of *Yersinia enterocolitica*. *Gene* **109**, 137-141 (1991).

14. Gerc AJ*, et al.* Visualization of the *Serratia* Type VI Secretion System Reveals Unprovoked Attacks and Dynamic Assembly. *Cell Rep* **12**, 2131-2142 (2015).

15. Shanks RM, Caiazza NC, Hinsa SM, Toutain CM, O'Toole GA. *Saccharomyces cerevisiae*-based molecular tool kit for manipulation of genes from Gram-negative bacteria. *Appl Environ Microbiol* **72**, 5027-5036 (2006).

16. Babin BM*, et al.* SutA is a bacterial transcription factor expressed during slow growth in *Pseudomonas aeruginosa*. *Proc Natl Acad Sci U S A* **113**, E597-605 (2016).

17. Choi KH, Schweizer HP. mini-Tn*7* insertion in bacteria with secondary, non-*glmS*-linked *att*Tn*7* sites: example *Proteus mirabilis* HI4320. *Nat Protoc* **1**, 170-178 (2006).

18. Meisner J, Goldberg JB. The *Escherichia coli rhaSR*-$P_{rhaBAD}$ Inducible Promoter System Allows Tightly Controlled Gene Expression over a Wide Range in *Pseudomonas aeruginosa*. *Appl Environ Microbiol* **82**, 6715-6727 (2016).

19. Cherepanov PP, Wackernagel W. Gene disruption in *Escherichia coli*: $Tc^R$ and $Km^R$ cassettes with the option of Flp-catalyzed excision of the antibiotic-resistance determinant. *Gene* **158**, 9-14 (1995).

20. Jacobs MA*, et al.* Comprehensive transposon mutant library of *Pseudomonas aeruginosa*. *Proc Natl Acad Sci U S A* **100**, 14339-14344 (2003).

21. Guzman LM, Belin D, Carson MJ, Beckwith J. Tight regulation, modulation, and high-level expression by vectors containing the arabinose $P_{BAD}$ promoter. *J Bacteriol* **177**, 4121-4130 (1995).

22. Rao VA, Shepherd SM, English G, Coulthurst SJ, Hunter WN. The structure of *Serratia marcescens* Lip, a membrane-bound component of the type VI secretion system. *Acta Crystallogr D Biol Crystallogr* **67**, 1065-1072 (2011).

23. Jack RL, Buchanan G, Dubini A, Hatzixanthis K, Palmer T, Sargent F. Coordinating assembly and export of complex bacterial proteins. *Embo J* **23**, 3962-3972 (2004).
